# Recurrent mutation in the ancestry of a rare variant

**DOI:** 10.1101/2022.08.18.504427

**Authors:** John Wakeley, Wai-Tong (Louis) Fan, Evan Koch, Shamil Sunyaev

## Abstract

Recurrent mutation produces multiple copies of the same allele which may be co-segregating in a population. Yet most analyses of allele-frequency or site-frequency spectra assume that all observed copies of an allele trace back to a single mutation. We develop a sampling theory for the number of latent mutations in the ancestry of a rare variant, specifically a variant observed in relatively small count in a large sample. Our results follow from the statistical independence of low-count mutations, which we show to hold for the standard neutral coalescent or diffusion model of population genetics as well as for more general coalescent trees. For populations of constant size, these counts are given by the Ewens sampling formula. We develop a Poisson sampling model for populations of varying size, and illustrate it using new results for site-frequency spectra in an exponentially growing population. We apply our model to a large data set of human SNPs and use it to explain dramatic differences in site-frequency spectra across the range of mutation rates in the human genome.

---

Recurrent mutation has long been recognized as an important factor of evolution (Fisher, 1928; Haldane, 1933; Wright, 1938). This is emphasized by recent analyses of single-nucleotide polymorphism (SNP) frequencies and variation of mutation rates across the human genome (Aggarwala and Voight, 2016; Harpak et al., 2016; Seplyarskiy et al., 2021) describing how patterns of variation depend on the mutation rate, particularly for rare variants. By a rare variant we mean an allele, such as an alternate base at a SNP, which is observed a relatively small number of times in a large sample. Unless the mutation rate is very small, indistinguishable copies of the same allele may descend from multiple mutations. Here we present a sampling theory for the numbers and associated frequencies of these unobserved or latent mutations in the ancestry of a rare variant.

Humans are on the low end of polymorphism levels among species (Leffler et al., 2012). On average, multiple mutations should be rare. In the 1000 Genomes Project data, about 1 in 1300 sites differ when two (haploid) genomes are compared, and SNPs with more than two bases segregating comprise only about 0.3% of the total SNPs observed (The 1000 Genomes Project Consortium, 2015). But polymorphism rates vary by two or three orders of magnitude depending on local sequence context (Aggarwala and Voight, 2016; Harpak et al., 2016; Seplyarskiy et al., 2021). Recurrent mutation is an important phenomenon for fast-mutating sites. Evidence for this can be found in the haplotype structure surrounding rare mutations (Johnson and Voight, 2020) and in the distribution of their frequencies among sites in large samples (Harpak et al., 2016; Seplyarskiy et al., 2021).

Here we focus on the latter, in particular on the site-frequency spectrum (Tajima, 1989; Braverman et al., 1995; Fu, 1995). Deviations in site-frequency spectra compared to standard predictions may be due to selection (Bustamante et al., 2001; Achaz, 2009; Ferretti et al., 2017), changes in population size over time (Eldon et al., 2015; Liu and Fu, 2015; Gao and Keinan, 2016) or population structure (Gutenkunst et al., 2009; Städler et al., 2009; Kern and Hey, 2017). But they may also be due to multiple mutations, i.e. to violations of the infinite-sites model assumption that each polymorphism is due to a unique mutation (Fisher, 1930a; Kimura, 1969, 1971; Ewens, 1974; Watterson, 1975).

The standard site-frequency prediction, which holds for a well-mixed population of constant large size *N* and neutral mutation rate *u* at a locus, is that the number of SNPs where a variant is found in *i* copies in a sample of size *n* should be proportional to *θ*/*i*, where *θ* = 4*Nu* (Tajima, 1989; Fu, 1995). This dramatically underpredicts the abundance of rare variants in data from humans, which is largely due to our recent explosive population growth (Keinan and Clark, 2012; Gazave et al., 2014; Gao and Keinan, 2016), but the standard neutral model is a useful starting point for modeling recurrent mutation.

Jenkins and Song (2011) studied the occurrence of one or two mutations at a single site under the standard neutral coalescent model (Kingman, 1982; Hudson, 1983; Tajima, 1983). They showed that if two mutations occur and are non-nested (meaning that all descendants of both mutations can be observed) there will be a shift away from rare variants and toward common ones. An earlier work focusing on the nested case is Hobolth and Wiuf (2009). Bhaskar et al. (2012) used a similar approach as Jenkins and Song (2011) to obtain results for one, two or three mutations, up to leading order in the mutation parameter *θ*. Sargsyan (2006, 2015) considered two mutations occurring at two different sites, and Jenkins et al. (2014) assume that two mutations are distinguishable and yield a tri-allelic polymorphism. These latter works (Sargsyan, 2006, 2015; Jenkins et al., 2014) allowed for variable population size following the general coalescent approach of Griffiths and Tavaré (1998). None of these works considered rare variants in particular but their predictions, especially those for non-nested mutations (Jenkins and Song, 2011; Bhaskar et al., 2012) are helpful for understanding recurrent mutation.

Two recent large studies of human SNPs observed this predicted shift away from rare variants and toward common ones at fast-mutating sites. Harpak et al. (2016) surveyed about 8 million SNPs in a sample of nearly 61 thousand people in version 0.2 of the Exome Aggregation Consortium database (Lek et al., 2016) for which data were available from other primate species. Among these, about 93.3% of these were bi-allelic, 6.5% were tri-allelic and 0.2% were quad-allelic. Harpak et al. (2016) took the presence of identical segregating variants in different species, ranging from chimpanzees to baboons, as indicative of a higher mutation rate at a site. Consistent with the hypothesis of multiple latent mutations at fast-mutating sites, they found fewer rare variants at bi-allelic SNPs for which the minor allele was segregating in another species, and that this effect is stronger when the other species is closer to humans.

The work we present here builds upon the second of these studies. Seplyarskiy et al. (2021) looked at rare variants in two datasets, one containing about 292 million variants among nearly 43 thousand individuals in TOPMed freeze 5 (Taliun et al., 2021) and the other containing about 182 million variants among 15 thousand individuals in gnomAD version r2.0.2 (Karczewski et al., 2020). Variants were divided into 192 types: each of the 3 possible base substitutions at the middle site of all 64 possible trinucleotides. A classic example of a fast-mutating site in this context would be ACG, which readily changes to ATG via a C to T transition at the CpG dinucleotide (Bird, 1980; Goldman, 1993). The main goals in Seplyarskiy et al. (2021) were to quantify how the rates of each kind of mutation vary across the genome and to partition this variation into distinct components correlated with different mutational processes.

Another aim, taken up in the Supplementary Materials of Seplyarskiy et al. (2021), was to correct for multiple mutations contributing to rare variants. Recurrent mutation was modeled as a multi-type Poisson process where mutations with lower sample counts occur independently at a locus to generate the appearance of higher count mutations (Desai and Plotkin, 2008). The expected counts in the absence of recurrence were taken from the site-frequency spectrum at slow-mutating sites. The loss of rare variants due to recurrent mutation at fast-mutating sites was quantified for sites with up to 70 copies of a rare variant. These were considered to have descended from up to 5 mutations. Slow-mutating sites, even with rates up to the genome average in humans, should conform fairly well to the infinite-sites assumption. Resampling from these as in Seplyarskiy et al. (2021) is a way of controlling for the myriad unknown factors affecting the site-frequency spectrum, including growth.

In this work, we present a sampling theory for latent mutations of rare variants at each given site-frequency count in a large sample. We describe a mathematical population genetic framework for the Poisson-resampling method in Seplyarskiy et al. (2021) and provide closed-form analytical expressions for several quantities of interest. We obtain new large-sample results for exponential growth and use these to illustrate the theory. We apply our results to a different subset of the gnomAD data than Seplyarskiy et al. (2021), synonymous variants observed in non-Finnish European individuals in v2.1.1, containing about 834 thousand variants at about 12.3 million sites among 57K individuals, presorted into 97 bins based on estimates of mutation rate by the method of Seplyarskiy et al. (2022, in prep.).

We develop and present these results in the next three sections. In Section 1, we begin with the standard neutral coalescent or diffusion model of population genetics (Ewens, 2004) and demonstrate a close connection between the Ewens sampling formula (Ewens, 1972) and distributions of latent mutations. In Section 2, we extend the results to populations which have changed in size, using the Poisson-sampling models of Watterson (1974a) and Arratia et al. (1992). In Section 3, we compare predictions for constant size to those for exponential growth and show how the new theory can be applied to understand the effects of recurrent mutation on counts of rare variants across the range of human per-site mutation rates.

## 1 Theory for constant-size large populations

In this section, we begin with a description of recurrent mutation via the well known predictions for allele frequencies in a population and in a sample at stationarity. We then use conditional ancestral processes to demonstrate independence of latent mutations of rare variants in a large sample, and show that their distribution is given by the Ewens sampling formula.

### 1.1 Stationary distributions and sampling probabilities

Consider a single locus with parent-independent mutation among *K* possible alleles in a population which obeys the Wright-Fisher diffusion (Fisher, 1930b; Wright, 1931; Ewens, 2004). Thus, the population is very large, well mixed, constant in size over time, and there is no selection. One unit of time in the diffusion process corresponds to 2*N_e_* generations (*N_e_* generations for haploid species) where *N_e_* is the effective population size. Each gene copy or genetic lineage experiences mutations at rate *θ*/2 and each mutation produces an allele of type *i* ∈ (1, …, *K*) with probability *π_i_*, with Σ*_i_ π_i_* = 1, independent of the allelic state of the parent. At stationarity, the joint distribution of the relative frequencies *x*_1_, …, *x*_*K*−1_ of alleles is given by

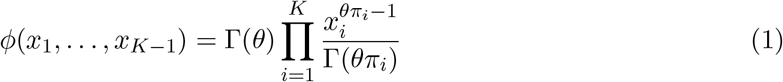

in which Γ(·) is the Gamma function, and where necessarily *x_K_* = 1 − Σ*_i<K_ x_i_* (Wright, 1931, 1949).

Conditional on the population frequencies (*X*_1_, …, *X_K_*) the sample counts of alleles 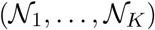 are multinomially distributed. A sample of size *n* taken from the population contains *n*_1_, …, *n*_*K*−1_ copies of alleles 1 though *K* − 1, and necessarily *n_K_* = *n* − Σ*_i<K_ n_i_* copies of allele *K*, with probability

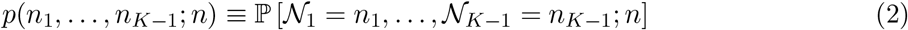

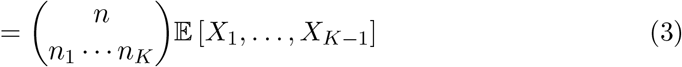

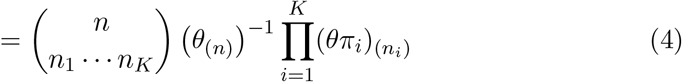

for *n_i_* ∈ (0, 1, …, *n*) constrained by Σ*_i_ n_i_* = *n* and where *k*_(*r*)_ denotes the Pochhammer function or rising factorial *k*(*k* + 1) · · · (*k* + *r* − 1) with *k*_(0)_ = 1. The shorthand defined in (2) is used extensively in what follows.

In applications to DNA, *K* = 4 and a sample at a given site would contain counts *n*_1_, *n*_2_, *n*_3_, *n*_4_ of each of the four nucleotides. The assumption of parent-independent mutation which leads to the relatively simple expressions (1) and (4) is unrealistic for DNA, but its results are useful in the case of rare variants in very large samples. In this case, it is likely that the common variant, allele 4 say, represents the ancestral state of the entire sample and that rare variants (alleles 1, 2 and 3) are due to recent mutations from the common variant. Then the mutation parameter *θπ_i_* for *i* ∈ (1, 2, 3) captures the production of type-*i* rare alleles in a specific ancestral background (allele 4).

An instructive special case is *K* = 2, where we have

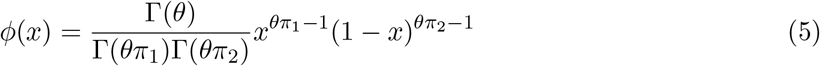

for the stationary distribution of the frequency of type 1 in the population Wright (1931), and

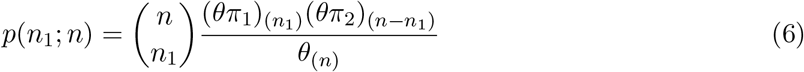

for the sampling probability, i.e. that a sample of size *n* contains *n*_1_ copies of allele 1 and *n*_2_ = *n* − *n*_1_ copies of allele 2. Any two-allele mutation model can be described as a parent-independent model, but this is not so in general for *K* > 2.

Figure 1 shows how the sample frequency distribution *p*(*n*_1_; *n*) in (6) depends on the mutation rate for a pair of alleles which differ by an order of magnitude in mutation rate. Three value of *θ* are shown (small, blue; middle, orange; large, red) with the small value chosen so that the mutation rate for allele 2 (*θπ*_2_) is equal to the human average of about 1/1300 and the mutation rate for allele 1 (*θπ*_1_) is ten times that. When *θ* is small, the distribution is U-shaped and nearly symmetric, given that the sample is polymorphic. When *θ* is around one, the distribution becomes J-shaped (or L-shaped if *π*_1_ < *π*_2_). When *θ* is large, the distribution has a peak around *π*_1_. Graphs of *ϕ*(*x*) (not shown) display these same shapes, and *p*(*n*_1_; *n*) will be very close to *ϕ*(*x*)*dx* when *n* is large.

**Figure 1:**
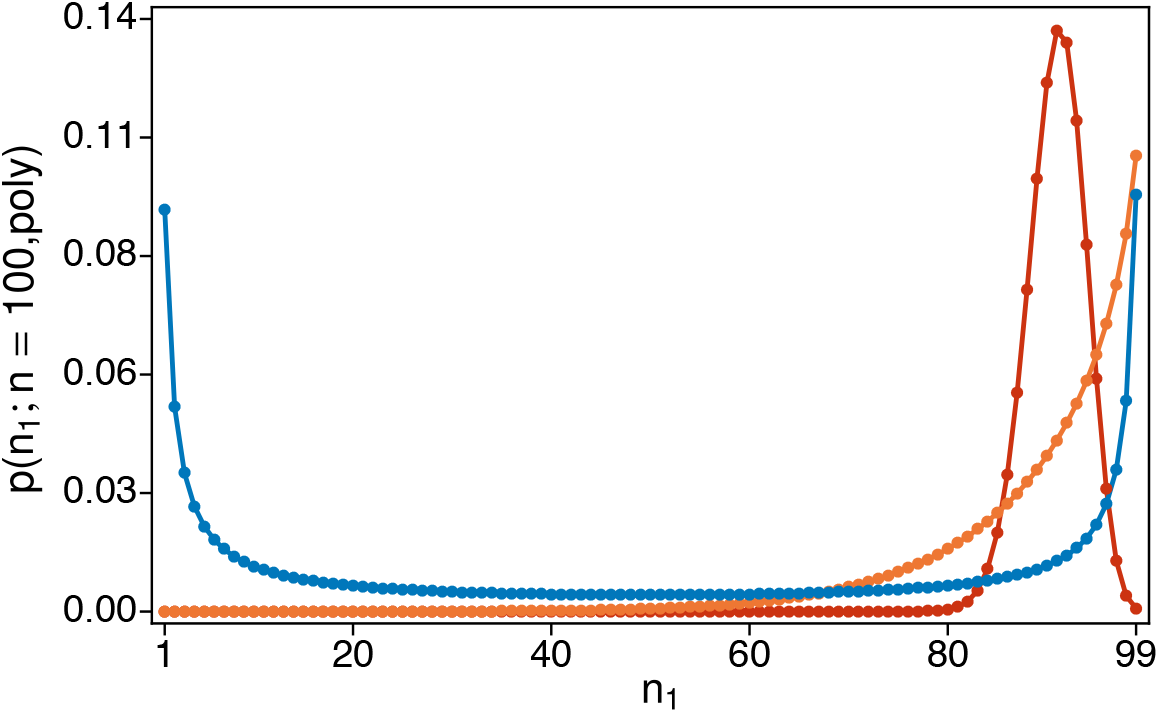
Sample frequency distribution *p*(*n*_1_; *n*) for *n* = 100, with *π*_1_ = 10*π*_2_ and three values of *θ* (smallest in blue, medium in orange, largest in red). The smallest *θ* was chosen so that *θπ*_2_ = 1/1300 ~ 0.00077, i.e. around the human average. The value of *θ* increases one-thousand fold from smallest to medium and again from medium to largest. In all three cases, the probabilities are normalized to sum to one, i.e. conditioned on the sample being polymorphic (1 ≤ *n*_1_ ≤ 99).

#### 1.1.1 Relationship to infinite-sites frequency spectra

We use *θ* for the per-site mutation parameter. In a collection of *L* total sites at which (6) holds, the finite-sites version of the site-frequency spectrum (i.e. the expected number of sites with *n*_1_ copies of allele 1 and *n*_2_ copies of allele 2) is given the product *Lp*(*n*_1_; *n*). Infinite-sites mutation models may be obtained as limits of finite-sites models as *L* tends to infinity with the total mutation parameter *Lθ* remaining finite. So when *θ* is small, we expect finite-sites results to be close to the usual (infinite-sites) predictions from the diffusion model (Ewens, 1979, 2004) or the coalescent model (Fu, 1995). Finite-sites models distinguish between kinds of mutations, subject to different mutation pressures, whereas infinite-sites models implicitly treat all mutations the same.

From Ewens (1979) equation (8.18) or Ewens (2004) equation (9.18)—see also Wright (1938) equation (16)—the expected number of sites segregating in the population with frequencies between *x* and *x* + *dx* under the infinite-sites model is proportional to 1/*x*. For comparison to (5) we may write

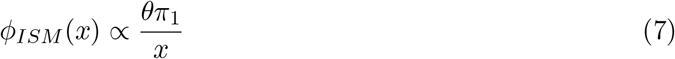

for a single site (*θ* small) approximately under the standard infinite-sites mutation model. For comparison with (6), we have

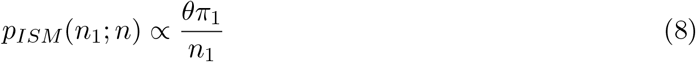

for the approximate single-site probability that there are *n*_1_ type-1 alleles in a sample of size *n*. Equation (8) has the same form as the usual infinite-sites site-frequency spectrum (Fu, 1995) but here it is for a specific mutant (allele 1) with a specific ancestral type (allele 2 in the two-allele model).

From (5) and (6) with *θ* small we have

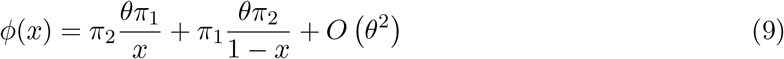

and

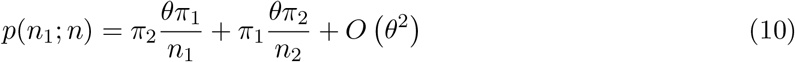

for *n*_1_ ∈ (1, …, *n* − 1). The diffusion result (5) does not admit atoms of probability at *x* = 0 or *x* = 1—see section 10.7 of Ewens (2004) for discussion—but we can interpret (9) intuitively as follows. If *θ* is close to zero, most of the time the population will be fixed, containing only allele 1 with probability *π*_1_ and only allele 2 with probability *π*_2_. Mutants of type 2 and type 1 are introduced with rates *θπ*_2_ and *θπ*_1_ in these two backgrounds, respectively. Then the leading terms in (9) represent a mixture of two infinite-sites models like (7) with the constants of proportionality specified. Equation (10) has an identical interpretation, as a mixture of two infinite-sites site-frequency spectra.

Although no closed-form expression like (1) is available except under parent-independent mutation, Burden and Tang (2016, 2017) have shown that the stationary densities for pairs of alleles under general mutation models take forms identical to (9) when *θ* is small; see equation (21) in Burden and Tang (2017). Similarly from a coalescent analysis of general *K*-alleles mutation, Bhaskar et al. (2012) obtained leading order terms for sampling probabilities with forms identical to (10) when *θ* is small and samples contain just two alleles. For *K* = 2, the result from Theorem 1 of Bhaskar et al. (2012) is identical to (10).

### 1.2 Mutation and the frequencies of rare sample variants

Our goal here is to understand how the frequency spectra of rare variants depend on *θ* and on the number of mutation events in the ancestry of the sample under the standard neutral coalescent or diffusion model of population genetics which assumes constant population size (Ewens, 2004). We first describe an ancestral process for the sample, then focus on rare variants in a large sample to obtain predictions about latent mutations.

#### 1.2.1 A conditional ancestral process for rare variants

Here we focus on ordered samples because the calculations are more intuitively related to the familiar rates of events in the ancestral coalescent process. The results do not depend on the order and so apply equally to ordered and unordered samples. Using the subscript “o” for ordered and writing *p*_o_(*n*_1_, …, *n_K_*) in place of *p*_o_(*n*_1_, …, *n*_*K*−1_; *n*) to facilitate the calculations, we have

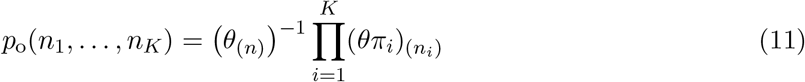

which differs from the sampling probability in (3) and (4) only by the multinomial coefficient, or the number of ways a sample containing allele counts *n*_1_, …, *n_K_* can be ordered.

Equation (11) is suggestive, as are (4) and (6), that the sampling structure of the *n_i_* copies of allele *i* may be related to the Ewens sampling formula (Ewens, 1972). Specifically, from the fact that

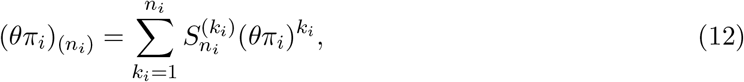

where 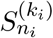 is an (unsigned) Stirling number of the first kind, we might guess that there is a latent variable *k_i_* which is the number of mutations giving rise to the *n_i_* copies of allele *i*. As in the usual application of the Ewens sampling formula, in contrast to the total possible number of type-*i* mutations in the ancestry of the sample, these latent mutations are just those *k_i_* ∈ (1, …, *n_i_*) most recent ones which produced the observed alleles.

That is, based on (11) and (12), we suppose that the joint probability of the sample counts *n*_1_, …, *n_K_* and their numbers of latent mutations *k*_1_, …, *k_K_* is given by

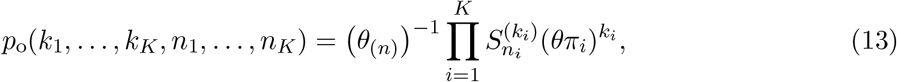

and therefore that the probability of *k*_1_, …, *k_K_* conditional on *n*_1_, …, *n_K_* is given by

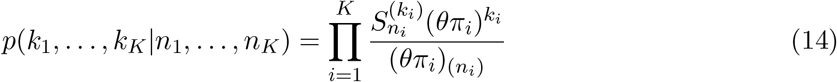

which applies to both ordered and unordered samples.

It is straightforward to check that (13) satisfies the corresponding recursive equation for the sampling probability, obtained using (17) in (15c) below and keeping track of latent mutations (not shown). It can also be obtained by the approach of Donnelly and Tavaré (1987), which begins with the Ewens sampling formula, labels mutations with allelic types 1 to *K* randomly (with probabilities *π*_1_ to *π_K_* in our notation) then retrieves the expected *K*-allele sampling probabilities.

Thus the number of latent mutations of an allele conditional on its sample count follows the Ewens sampling formula. But reckoned in this way under parent-independent mutation some of the latent mutations in (13) and (14) are ‘empty’ (Baake and Bialowons, 2008). They do not change the allelic type. These are a modeling artefact which must be dealt with not only in parent-independent models but in general mutation models as well the way these are typically implemented (Jenkins and Song, 2011; Bhaskar et al., 2012; Jenkins et al., 2014; Burden and Tang, 2017; Burden and Griffiths, 2019). Empty mutations have no empirical significance. Here we show that they almost never occur in the ancestry of rare variants in large samples.

We make use of the ancestral-process approach developed by Griffiths and Tavaré (1994a,b) based on recursive equations for sampling probabilities. For the *K*-allele model we have

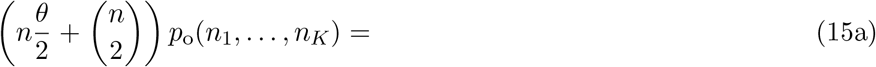

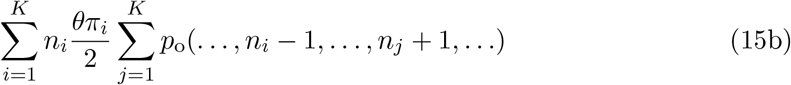

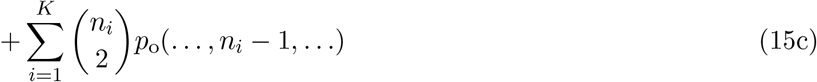

with boundary conditions *p*_o_(0, …, *n_i_* = 1, …, 0) = *π_i_* for *i* ∈ (1, …, *K*). This is a recursion back into the ancestry of the sample, in which (15b) and (15c) include all events which could have produced the sample, and the probabilities of ancestral types required in each case to produce the sample. It can be derived either a using coalescent approach (De Iorio and Griffiths, 2004) or a diffusion approach (Burden and Griffiths, 2019). We have kept the common factors of 1/2 in (15a), (15b) and (15c) to emphasize that the ancestral process occurs on the diffusion or coalescent time scale, with mutations happening at rate *θ*/2 on each ancestral lineage and coalescent events happening at rate 1 for each pair of ancestral lineages.

Initially these ancestral processes were used to compute otherwise intractable likelihoods analytically for small samples or by simulation for large samples (Griffiths and Tavaré, 1994a,b). They were subsequently used to describe the joint sampling of gene genealogies and associated allelic types under selection (Krone and Neuhauser, 1997; Neuhauser and Krone, 1997), and, in cases were the sampling probabilities are known, to describe conditional ancestral processes of samples given their allelic types (Slade, 2000a,b; Fearnhead, 2002; Stephens and Donnelly, 2003; Baake and Bialowons, 2008). We take the latter approach to obtain a large-*n* approximation for the conditional ancestral processes of mutation and coalescence for samples containing mostly one type. We set *K* to be the common allele, so we will have *n_K_* ⨠ *n*_1_, …, *n*_*K*−1_ and *n_K_* ~ *n*.

Equations (15b) and (15c) include, respectively, all possible mutation events and all possible coalescent events in the ancestral process which have a non-zero chance of producing the sample (*n*_1_, …, *n_K_*). Note that the terms with *i* = *j* in (15b) are the ‘empty’ mutations of Baake and Bialowons (2008).

Following Slade (2000b), the conditional ancestral process remains in state (*n*_1_, …, *n_K_*) for an exponentially distributed time, i.e. leaving that state at rate *n*(*θ* + *n* − 1)/2, then jumps to each possible ancestral state with probabilities proportional to the terms in (15b) and (15c). Dividing through by the left-hand side, (15a), we have

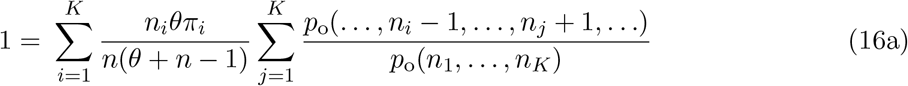

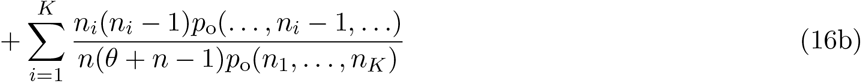

for the total probability of these events given that an event occurs in the ancestral process. These are, in (16a), mutations on lineages ancestral to alleles of type *i* which have alleles of type *j* as ancestors and, in (16b), coalescent events between lineages ancestral to alleles of type *i*.

Only the numbers ancestral lineages are tracked in (16a) and (16b). The full ancestry, or gene genealogy, can be modeled using exchangeability within each allelic type. That is, each of the *n_i_*(*n_i_* − 1)/2 pairs is equally likely to be involved in a type-*i* coalescent event and each of the *n_i_* lineages is equally likely to be the one involved in a type-*i* mutation event.

Depending on what quantities or aspects of the ancestry are of interest, (16a) and (16b) may be augmented, simplified or otherwise rearranged. Here we follow Fearnhead (2002) and Baake and Bialowons (2008), in removing some lineages from the ancestral process once they have experienced a mutation. This is captured by the identity

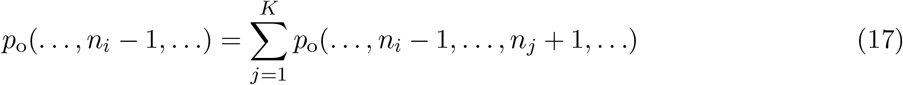

which can be used as needed in (16a). Our aim here is to model mutation and coalescence in the ancestry of the rare alleles with counts *n*_1_ through *n*_*K*−1_ in the sample. So we follow the ancestry of the *n_K_* common alleles only insofar as this affects the ancestries of the rare alleles. We use (17) to justify removing ancestral type-*K* lineages whenever they mutate, and we lump these events with coalescent events because their overall effect is the same (*n_K_* → *n_K_* − 1). Additionally, we distinguish two kinds of mutation events among the rare alleles: ones in which the ancestral allele was the common allele *K* and ones in which it was a rare allele *j* ∈ (1, …, *K* − 1).

Making these changes, and using (11) to simplify the ratios of sampling probabilities, we have

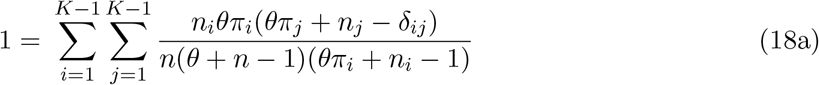

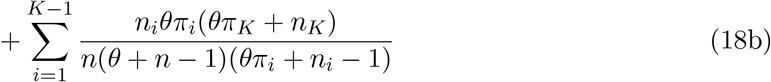

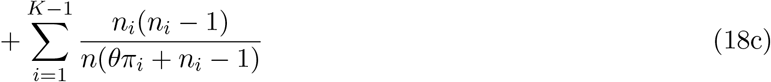

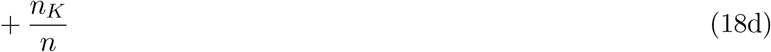

in which we have used Kronecker’s delta to accommodate empty mutations, *i* = *j* in (18a). Recall that *n* = Σ*_i_ n_i_* which will be *O*(*n_K_*) when *n_K_* becomes large for given *n*_1_ though *n*_*K*−1_. Equation (18a) contains the probabilities of all mutations on rare-allele lineages which have rare-allele ancestors. These probabilities are 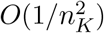 when *n_K_* is large. Equation (18b) contains the probabilities of all mutation events on rare-allele lineages which have common-allele ancestors. These are *O*(1/*n_K_*) when *n_K_* is large. Equation (18c) contains the probabilities of all coalescent events between rare-allele lineages of the same type, similarly *O*(1/*n_K_*). Finally, (18d) gives the probability of mutation or coalescence among the common-allele lineages, which is *O*(1).

Keeping only up to the *O*(1/*n_K_*) terms gives an approximate, large-*n_K_* ancestral process with total rate 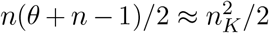 and jumps, for *i* ∈ (1, …, *K* − 1), from state (*n*_1_, …, *n_K_*) to state

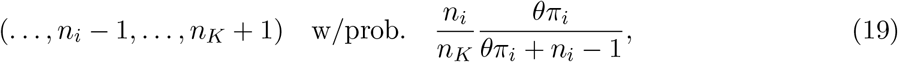

to state

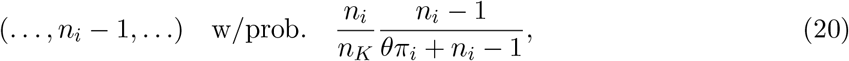

or to state

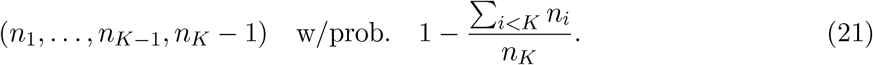

This process is dominated by (21), that is by events on lineages ancestral to the common allele *K*, which decrease the number of these but leave the counts of rare-allele lineages unchanged. Although we are not tracing the details of common-allele ancestry, we note that the overwhelming majority of these events will be coalescent events, since their rate is approximately equal to the total rate 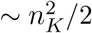. The next most frequent will be empty mutation events at rate *O*(*n_K_*), followed by common-allele mutation events with rare-allele ancestors at rate *O*(1).

When one of the rarer events occurs in the ancestral process, it involves allele *i* with probability *n_i_/n_K_*, then is either a mutation event from a common allele as in (19) or a coalescent event as in (20). For each allele *i* ∈ (1, …, *K* − 1) which is observed at least once in the sample, there will be exactly *n_i_* such events. Again, the empty mutation events captured in (18a) are negligible for large *n_K_*. Note that, the relative probabilities of mutation versus coalescence in (19) and (20) for each allele are identical to the standard ones from coalescent theory (Kingman, 1982), only here with *θπ_i_* in place of the usual *θ*. It follows that both the number of latent mutations which produced the *n_i_* copies and the counts of each mutation’s descendants among the *n_i_* copies are given by the Ewens sampling formula (Ewens, 1972; Kingman, 1982; Arratia et al., 1992, 2016).

The events involving the common allele in (21) occur very quickly. But since only a fixed number of events involving rare alleles are required to resolve the ancestry of latent mutation and coalescence, the approximation remains accurate until all the rare-allele events have happened, if *n_K_* is large enough. In Appendix section A.1, we study the joint distribution of the times of events among the rare alleles and the numbers of common-allele ancestors when these rare-allele events occur. Focusing on the case of two alleles for simplicity, if 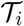 is the time back to the *i*th event involving the rare allele 1, we have

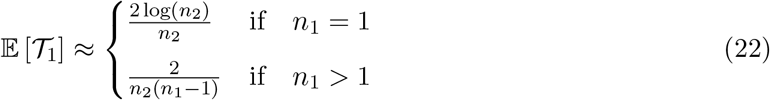

which in either case tends to zero as *n*_2_ tends to infinity. Further, if 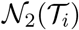 is the random number of type-2 ancestral lineages left at the *i*th event involving the rare allele 1, we have

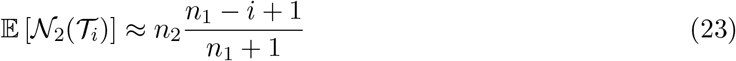

suggesting that, despite the rapid decrease of common-variant lineages, the approximation can hold until the entire ancestry of latent mutation and coalescence is resolved.

Even for the largest rare-variant site-frequency count considered in Seplyarskiy et al. (2021), there will still be > 1200 common-variant lineages left on average at 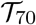 for the TOPMed data (*n*_2_ ~ 86000) and > 400 left for the gnomAD data (*n*_2_ ~ 30000). In Section 3.2, we consider site-frequency counts up to 40 for synonymous exonic sites in gnomAD with many fewer SNPs but a larger sample size (*n*_2_ ~ 114000) and in this case there should be about 2780 common-variant lineages left at 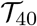 when the entire ancestry of latent mutation and coalescence among the rare variants is resolved.

Thus, rare alleles in a large sample will quickly coalesce and mutate. Their ancestors will be common alleles. If *k_i_* ∈ (1, …, *n_i_*) is the number of mutations in the ancestry of allele *i* ∈ (1, …, *K* − 1), then from the rates of mutation and coalescence in (19) and (20) we have

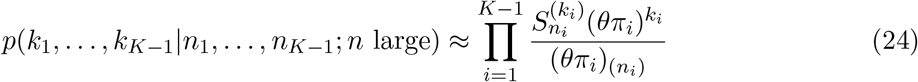

which is the product of independent Ewens distributions.

#### 1.2.2 Latent mutations and sample counts of rare alleles

Here we focus on the case in which a single type of mutation or allele is observed against a background of a given common allele, as in recent empirical studies (Harpak et al., 2016; Seplyarskiy et al., 2021). Our goal is to understand how counts of these mutant alleles depend on the number of latent mutations and on the mutation rate. As before, allele 1 is the focal rare allele and allele *K* is the common allele.

First, from (24) we have

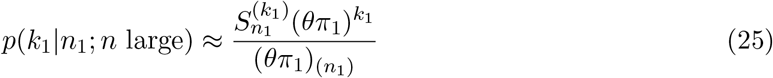

which sums to one for *k*_1_ ∈ (1, …, *n*_1_). To understand the effects of mutation on the number of rare alleles, we apply Stirling’s formula as in 6.1.47 of Abramowitz and Stegun (1964) or equation (1) in Tricomi and Erdélyi (1951) to show that the main dependence of *p*(*n*_1_; *n*) on *n*_1_ and *θπ*_1_ is captured by

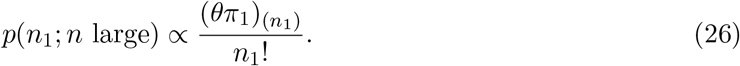

Using this together with (25) we have

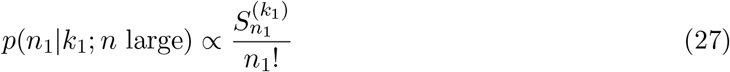

as the approximate dependence of the rare-allele count *n*_1_ on the number of latent mutations *k*_1_. Following the logic of Section 1.1.1, we can use (26) and (27) to understand how the site-frequency counts of a rare allele depend on the rate of production of the allele and on the number of latent mutations contributing to those counts.

The proportional relations (26) and (27) are sufficient for this if we adopt the usual convention of normalizing site-frequency counts to sum to one. Figure 2A shows how the site-frequency spectrum of a rare variant in a large sample depends on the number of mutations which produced the observed copies of the variant. When all copies descend from a single mutation (*k*_1_ = 1), the usual predictions from the infinite-sites model hold. Thus, putting 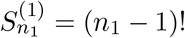 in (27) we have

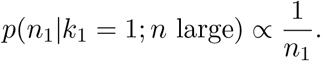

The total number of such sites will depend on *θπ*_1_, and in general on the factor 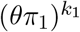 in (25) for larger numbers of latent mutations. But conditional on *k*_1_, the normalized site-frequency counts for a rare variant do not depend on *θ*, at least to leading order in the sample size *n* ~ *n_K_*. Further, if there are *k*_1_ > 1 mutations in the ancestry of the rare variant, then *n*_1_ cannot be less than *k*_1_. This is illustrated in Figure 2A for *k*_1_ = 2 to *k*_1_ = 5 latent mutations. A key effect of recurrent mutation is to give relatively less weight to low site-frequency counts, as found previously by Jenkins and Song (2011).

**Figure 2:**
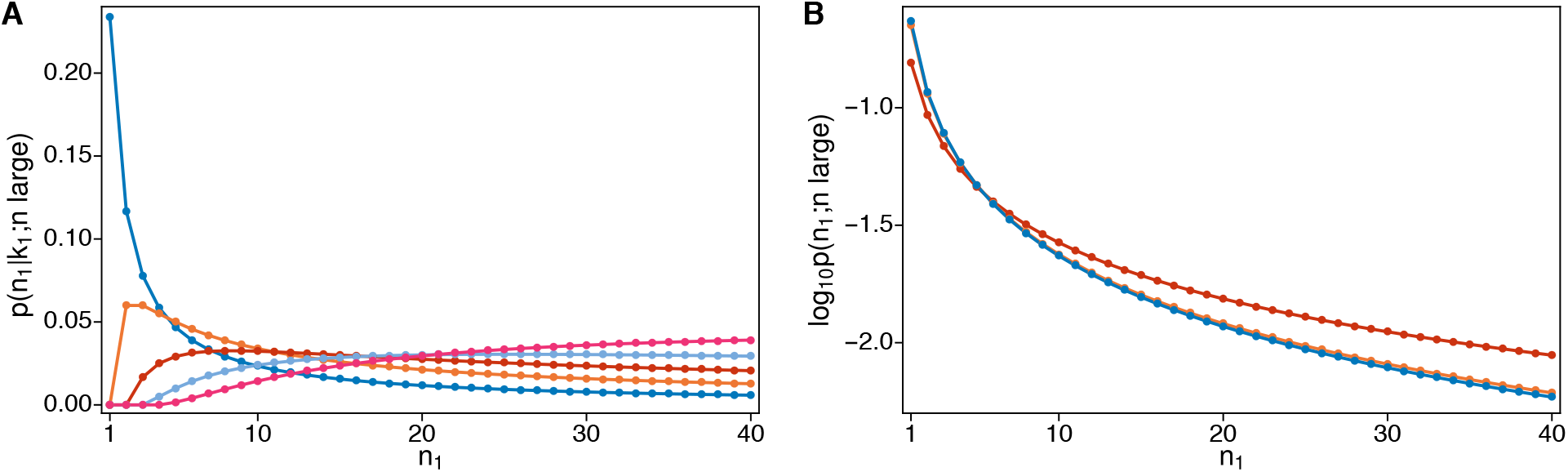
Panel A shows the probability of observing *n*_1_ copies of allele 1 in a large sample given these are produced by *k*_1_ = 1, 2, 3, 4, 5 mutations. Panel B shows the log_10_-probability of observing *n*_1_ copies of allele 1 in a large sample for three different values of *θπ*_1_: 0.002, 0.02 and 0.2. Probabilities in both panels are normalized to sum to one for *n*_1_ ∈ (1, 2, …, 40).

Using (25) and (26) the joint distribution of *n*_1_ and *k*_1_ obeys

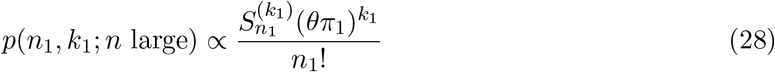

which can be compared to the results of Jenkins and Song (2011). With fixed *n*_1_ and large *n_K_* in our model, all mutations in the ancestry of the rare variant will be non-nested mutations; note this also follows from (18) in Jenkins and Song (2011). In addition, we have shown that the higher-order terms in *θ*, i.e. beyond 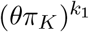, can be neglected when *n_K_* is large. Adapting the notation of Jenkins and Song (2011) in which 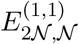 is the event that the *n*_1_ copies of allele 1 are due to two non-nested mutations, both from allele *K* to allele 1, their (21) becomes

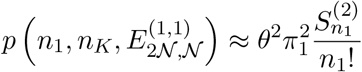

for large *n_K_* (and small *θ*), which is identical to (28) if *k*_1_ = 2.

Numerical computations (not shown) using the unnumbered equation below (10) in Jenkins and Song (2011), which holds for any *θ*, reproduce the case of *k*_1_ = 2 shown in Figure 2A when *n_K_* is large. This is evident in Figure 3 of Jenkins and Song (2011) for the quantity 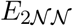. These computations are difficult for samples beyond the hundreds. Our results for *k*_1_ = 3 could potentially also be compared to the *O*(*θ*^3^) results of Bhaskar et al. (2012) using their Theorem 3 and summing appropriately.

**Figure 3:**
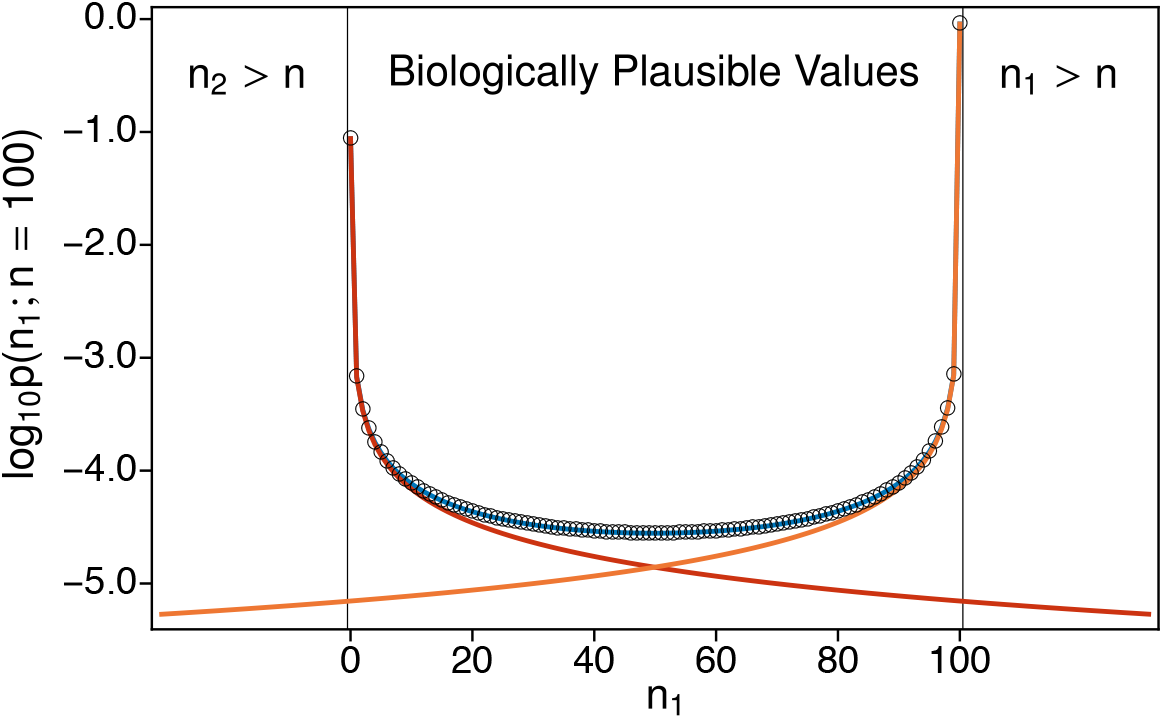
Schematic relating the independent-Poissons model to the classic two-allele diffusion result (6). Parameters are as in Figure 1, specifcally *θπ*_2_ = 1/1300, *π*_1_ = 10*π*_2_ and *n* = 100. The blue line shows the sum (42). The red line shows the first term of that sum. The orange line shows the second term. The plot extends beyond the biologically plausible values of *n*_1_ because the independent-Poissons model allows *n*_1_ > *n* and *n*_2_ > *n*. Open circles show (6) which closely tracks the blue line across the range of biologically plausible values, and the red and orange lines for small *n*_1_ and small *n*_2_ respectively. Biologically implausible are rare in this case.

Figure 2B shows how the site-frequency counts of the rare variant depend on the mutation parameter of that variant, *θπ*_1_. Although Figure 2A shows a dramatic effect of *k*_1_ on the site-frequency counts, Figure 2B suggests that large values of *k*_1_ are unlikely. This is evident from (25) and (28) in that each additional mutation results in an additional factor of *θπ*_1_. Note that the smallest value of *θπ*_1_ in Figure 2B is already more than twice the human average. From (26), we have

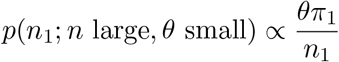

which is consistent with (10) in the case where allele 1 is rare in a large sample. Thus, when *θπ*_1_ is small (0.002 and 0.02 in Figure 2B) the site-frequency spectrum under recurrent mutation is very close to the standard infinite-sites model predictions. When *θπ*_1_ is large (0.2 in Figure 2B) the site-frequency spectrum under recurrent mutation is noticably different, with a dearth of low-frequency variants and corresponding excesses at higher frequencies. Figure 2B plots site frequencies on a log scale to better illustrate differences, especially at higher frequencies.

## 2 Theory for nonconstant populations

Here we extend our analysis to populations which deviate from the standard neutral site-frequency predictions. We have in mind populations which have changed in size, although other applications may be possible. Now gene genealogies are the general coalescent trees of Griffiths and Tavaré (1998), which have same the branching structure of standard coalescent trees but may have different distributions of coalescence times.

Equation (25) suggests another way to model both the number of copies (*n*_1_) of a variant of interest and the corresponding count of latent mutations (*k*_1_) when the variant is rare in a large sample. Arratia et al. (1992) proved that when the sample size tends to infinity, the numbers of alleles in small counts 1, 2, …, *i* in the Ewens distribution converge to independent Poisson random variables with expected values *θ, θ*/2, …, *θ/i*. Note that *θ/i* is the usual expected site-frequency count of mutants in *i* copies in the sample under the standard neutral model of a large constant-size population. A seminal result of Watterson (1974a) is that the numbers and counts of mutations in a sample from such a multi-type Poisson distribution conform to the Ewens sampling formula when conditioned on their total size. So we may interpret (25) and other findings in the previous section within this independent-Poissons sampling framework.

This is exactly the approach in the Supplementary Materials of Seplyarskiy et al. (2021). Again, human SNP data strongly reject the standard neutral model with site-frequencies ∝ 1/*i*, owing largely to the great excess of singletons and other rare variants due to our recent growth (Keinan and Clark, 2012; Gazave et al., 2014). So we replace 1/*i* with 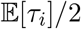, where *τ_i_* is the total length of branches with *i* descendants in the gene genealogy of a sample. For an extension of independent-Poissons sampling to variants under selection, see Desai and Plotkin (2008). Our notation is different than in Seplyarskiy et al. (2021) because here we use the coalescent or diffusion time scale.

Under the standard neutral coalescent model, 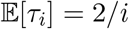. For the general coalescent trees of Griffiths and Tavaré (1998), *τ_i_* can be expressed in terms of the coalescent intervals, *T_k_*, which are the lengths of time when there were *k* ∈ (2, …, *n*) lineages in the ancestry of the sample. In particular,

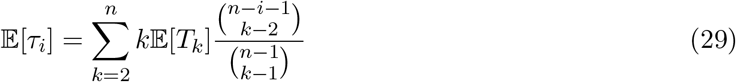

(Fu, 1995; Griffiths and Tavaré, 1998).

Watterson (1974a) studied three models. In Model 1, using our notation, mutations arise from a constant source at rate *θ*, then propagate or go extinct independently according to a critical branching process, i.e. with birth rate equal to death rate as for a neutral mutation. The number of mutations in count *i* has expected value *θμ^i^/i*, for a constant *μ* > 0 which converges to 1 as the duration of the process increases. Watterson (1974a) proved that the numbers and counts of mutations follow the Ewens sampling formula when conditioned on their total size, which for Watterson (1974a) was equivalent to the population size. Models 2 and 3 are the Moran model and the Wright-Fisher model (Moran, 1958, 1962; Fisher, 1930b; Wright, 1931) and Watterson (1974a) proved that these have the same limit as Model 1 when the population size is large.

Model 1 is an example of a logarithmic species distribution (Fisher, 1943; Watterson, 1974b; Arratia et al., 2003; Lambert, 2011). Branching-processes have also been used to describe and infer the ages of rare alleles (Rannala and Slatkin, 1997; Slatkin and Rannala, 2000; Wiuf, 2000); for recent developments and a review, see Crespo et al. (2021). Slatkin (2000) used this approach and an extension of Griffiths and Tavaré (1998) to model the ages of rare alleles in a large sample. Champagnat and Lambert (2012, 2013) studied the convergence of population frequencies of alleles for supercritical, subcritical or critical branching processes. All of these works assume that each allele traces back to a single mutation, as under the infinite-alleles mutation model.

Our approach to modeling recurrent mutation follows that of Watterson (1974a) to Model 1. Whereas Watterson (1974a) did not specify the source of mutations, here we take it to be the production of rare variants by mutation from a common variant on the gene genealogy of a large sample. What for Watterson (1974a) was the total population size is for us the total count of a rare variant. Allele 1 is our nominal variant of interest, but for simplicity for the moment, we use *n*, *k* and *θ* in place of *n*_1_, *k*_1_ and *θπ*_1_. As a further notational convenience, we define

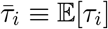

so that 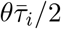 is the expected number of mutations with count *i* in this independent-Poissons sampling model.

Let (*a*_1_, *a*_2_, …) be the numbers of latent mutations of the variant of interest with counts (1, 2, …). We assume that 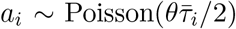 and that *a_i_* and *a_j_* are independent for *i* ≠ *j*. Their joint distribution is then

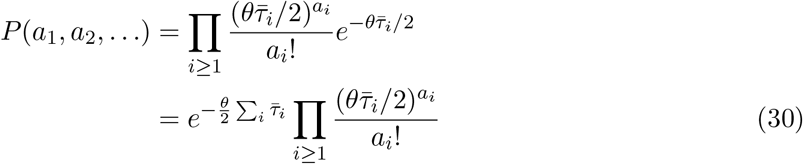

with *a_i_* ≥ 0. The total sample size is what would set the upper limits of the product and the sum above, but we leave these unspecified for now, only imagining that the total sample size is much larger than the sample count of the variant of interest, so we can model the latter without restriction.

We are only concerned with *a_i_* for *i* ≤ *b*, where *b* is the largest rare-variant count. Thus, the assumption of independence in (30), which is equivalent to there being no nested mutations in the ancestry of a rare variant, will only need to be true for 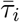 with *i* ∈ (1, …, *b*). In Appendix section A.2 we prove that this holds for the trees of Griffiths and Tavaré (1998) for fixed *b* in the limit as the total sample size tends to infinity, and that the counts (*a*_1_, …, *a_b_*) converge to independent Poisson random variables as with expected values 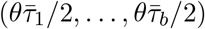. A condition is that the total height of the genealogy is bounded, which is a mild assumption ruling out pathological situations such as a populations whose sizes *increase* too quickly backward in time.

The count of the variant of interest is *n* = Σ*_i_ ia_i_* and its number of latent mutations is *k* = Σ*_i_ a_i_*. Following Watterson (1974a), we consider the probability generating function of *n* and *k*, which in the present case simplifies to

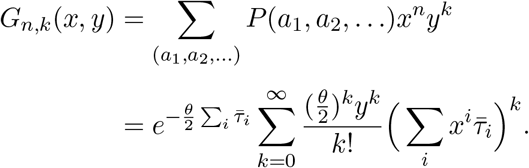

For the details of this derivation, see (82) in the Appendix. The coefficient of *x^n^* (and *y^k^*) can be found using

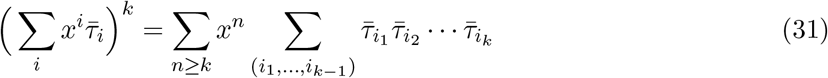

where the sum is over

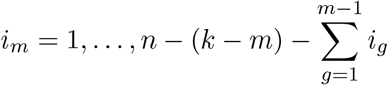

for *m* = 1, …, *k* − 1, and with

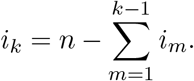

Returning to our notation in which *n*_1_ is the number of copies of a variant of interest, *k*_1_ its number of latent mutations, *θπ*_1_ its mutation parameter, and *n* is the total sample size, and further using *τ* to show the new dependence on the vector of expected times 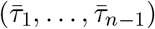, we have

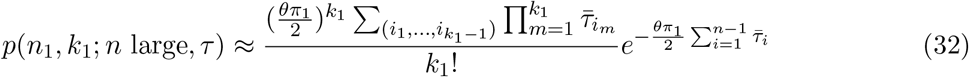

which is non-zero for *n*_1_ = *k*_1_ = 0 and *n*_1_ ≥ *k*_1_ ≥ 1. The sum over 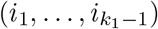 here is the same as in (31). It is equivalent to summing over partitions of the integers 1 through *n*_1_ into *k*_1_ subsets, where the sizes of the subsets are 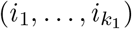.

It is convenient to decompose (32) as follows. The number of type-1 mutations is Poisson distributed

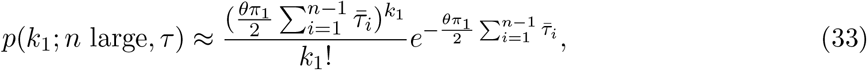

with parameter equal to the expected number of type-1 mutations on the gene genealogy of the sample. Conditional on this, the distribution of the number of times allele 1 appears in the sample is given by

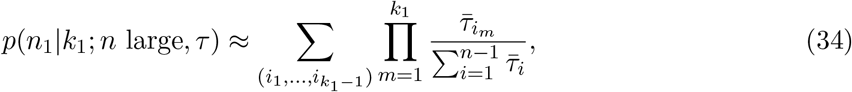

which depends on the relative expected branch lengths but does not depend on *θ* or *π*_1_.

Alternatively, *p*(*n*_1_; *n* large, *τ*) can be computed by summing (32) appropriately, over *k*_1_ ∈ (0, …, *n*_1_). Then

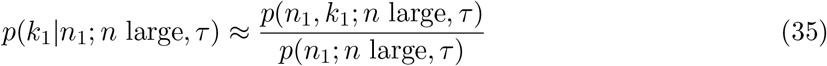

can be used to estimate the number of independent mutations which produced the observed copies a rare allele.

The sum over 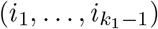 in (34) and (32) is straightforward to evaluate but will become impractical if *n*_1_ and *k*_1_ become too large. In what follows, we consider *k*_1_ ≤ 7 mutations at a each site. Equation (33) suggests that this will be accurate up to about three expected mutations per site, because the probability of *k*_1_ greater than 7 is just over 1% when 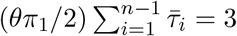. As in Figure 2, the largest value of *n*_1_ we consider is 40. These are not the upper limits of feasibility; it takes two minutes to evaluate (34) for all *k*_1_ ∈ (0, …, 7) and *n*_1_ ∈ (0, …, 40) in Mathematica version 11.2 (Wolfram Research, Inc., 2017) on a mid-2015 MacBook Pro.

Considering the first three possible values of *k*_1_ in (34),

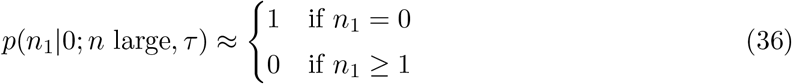

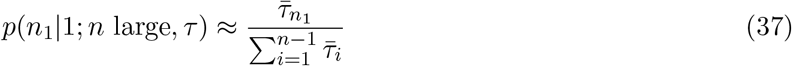

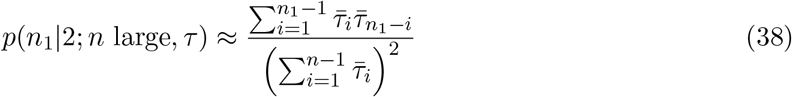

Equation (36) says simply that if there are no type-1 mutations on the gene genealogy then no copies of allele-1 will be observed. Equation (37) is the familar result for the site-frequency spectrum, that it is given by the proportion of branches in the tree that have *n*_1_ descendants. Equation (38) extends this to two mutations and emphasizes that mutations in the ancestry of a rare allele will be non-nested when *n* is large.

For the constant-size model, we can put 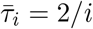 in (32) and obtain new expressions corresponding to (26), (27) and (28),

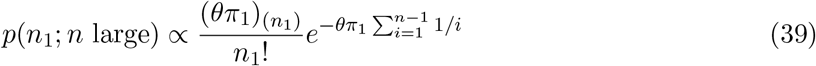

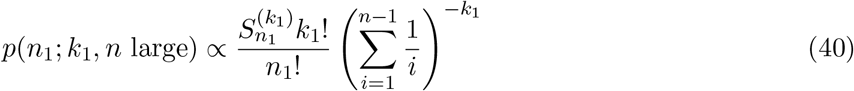

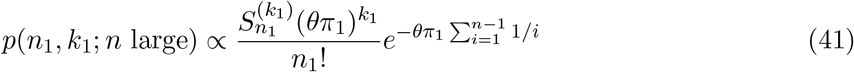

which may be preferable to the previous ones. The expression for *p*(*k*_1_|*n*_1_; *n* large) obtained using 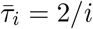 here is the identical to (25). Figure 2 is also unchanged if (39) and (40) are used instead of (26) and (27), and normalizing in the same way.

### 2.1 Relation to *K*-alleles diffusion results

These new results may be discerned in the sampling probabilities from the diffusion model. For example, a more detailed treatment of (6) and application of Stirling’s formula gives

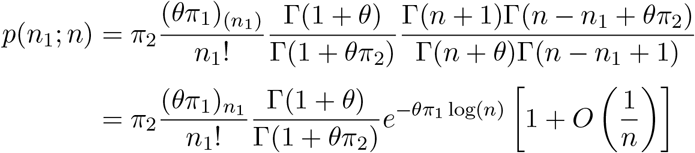

in which we write 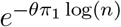 in place of 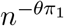 to emphasize the connection to the gene genealogy. Using a Taylor series approximation for the Gamma function around 1 we have

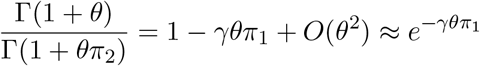

where *γ* = 0.577 ‥. is Euler’s constant. As 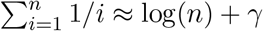 when *n* is large, we find that (6) will be very close to *π*_2_ times the right-hand side of (39) for a given *n*_1_ when *n* is large and *θ* is small. For reference, note that even the fastest-rate sites in the human genome have *θ* only equal to about 0.02 (see Section 3.2 below).

In deriving (32), we assumed that both the number of mutations and their total count (*k*_1_ are *n*_1_ here) are unbounded. We did not take a formal limit as *n* → ∞, and instead have 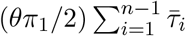 for the rate of occurence of type-1 mutations. This is intuitive from the standpoint of coalescent theory, since 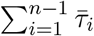 is the average total length of the gene genealogy, but allowing that *k*_1_ are *n*_1_ could potentially be larger than *n* makes little sense. These new results (32) through (41) are for rare alleles only, that is for a given *n*_1_ when *n* is very large.

To relate these new results to the model in Section 1.1, we can follow the logic in Section 1.1.1 and approximate the full sampling probability (6) when *θ* is small as the sum of two copies of the independent-Poissons sampling process. Thus, similar to (10), for the two-allele case here we can approximate the sampling probability *p*(*n*_1_; *n*) in (6) using

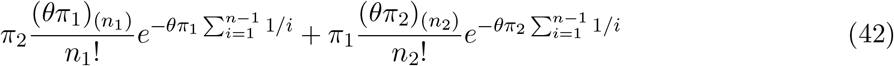

in which we have used result (39) twice, once for type-1 mutations in a type-2 ancestral background and once for type-2 mutations in an type-1 ancestral background, where it is understood that *π*_2_ = 1 − *π*_1_ and *n*_2_ = *n* − *n*_1_. This can be seen as a sample-based version of the boundary mutation model (Vogl and Clemente, 2012; Schrempf and Hobolth, 2017; Vogl et al., 2020) but one which allows for multiple segregating mutations.

Figure 3 plots the two terms in (42) individually, together with their sum (42) and *p*(*n*_1_; *n*) from (6). The parameters as the same as the small-*θ* case in Figure 1, specifically a sample of size *n* = 100 with *θ* chosen so that the mutation rate for allele 2 (*θπ*_2_) is equal to the human average (1/1300) and *π*_1_ = 10*π*_2_. In contrast to Figure 1, the range of *n*_1_ (similarly *n*_2_) in Figure 3 extends beyond what is biologically plausible. For these parameters, the expected number of type-1 mutations on the gene genealogy is about 0.04 and the expected number of type-2 mutations is about 0.004. Only about 1/50 polymorphic sites where the rare variant is allele 1 are expected to have experienced more than one mutation, and the corresponding value for sites where the rare variant is allele 2 is about 1/500. The different probabilites at the boundaries 0 and 100 result both from the ten-fold greater rate of type-1 versus type-2 mutation on the gene genealogy and the weights *π*_2_ and *π*_1_ in (42) which similarly capture the ten-fold difference in the the chance of the ancestral allele being of type 2 or of type 1, respectively, at stationarity in the Wright-Fisher diffusion model.

## 3 Theoretical example and data application

Here we illustrate the theoretical and empirical use of (33) and (34). First we describe the consequences of recurrent mutation in an exponentially growing population compared to those in a population of constant size. Second we explore an entirely empirical application to human SNP data, which suggests that disparate site-frequency spectra may be explained by differences in mutation rate (and thus recurrent mutation).

Note that if estimates of the expected fraction of the gene genealogy comprised of branches with *i* descendants, that is

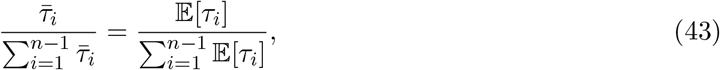

are available, then *p*(*n*_1_|*k*_1_; *n* large, *τ*) can be computed using (34). In addition, for any estimated or supposed values of the expected number of mutations on the gene genealogy,

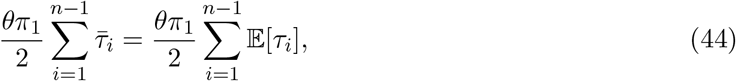

the joint distribution of the number of latent mutations, *k*_1_, and their total count, *n*_1_, is the product of (33) and (34).

### 3.1 An exponentially growing population

Consider the simple model of pure exponential growth which has been the subject of a number of studies (Slatkin and Hudson, 1991; Griffiths and Tavaré, 1998; Polanski and Kimmel, 2003; Chen and Chen, 2013; Polanski et al., 2017): a population which has reached its current (haploid) size *N*_0_ by exponential growth at rate *r* per generation. On the coalescent time scale of *N*_0_ generations, looking backward in time and setting *α* = *N*_0_*r*,

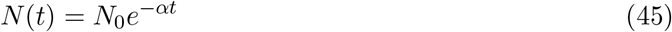

gives the population size at time *t* in the past. This model is unrealistic because the past population size approaches zero, but it can be taken as a rough approximation for recent dramatic growth. For instance, a population of current size *N*_0_ = 5×10^7^ with a generation time is 30 years and *r* = 0.0064, would have *α* = 3.2 × 10^5^. About 40, 000 years ago, it would have had size 10^5^, and using equation (7) in Slatkin and Hudson (1991) the pairwise coalescence time would be about 57, 000 years.

The expectation 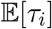 can be computed from (29) if the expected coalescent intervals 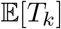 are known. We use the large-*n* results of Chen and Chen (2013) for 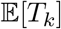 (our notation) to obtain a simple approximation for 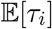. With the time scale and notation here, equation (11) in Chen and Chen (2013) gives

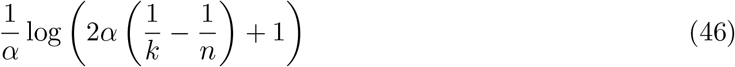

as a large-*n* approximation for the cumulative expected time for the number of ancestral lineages of the sample to decrease from *n* to *k*. Writing (46) as a continuous function of *x* = *k/n*,

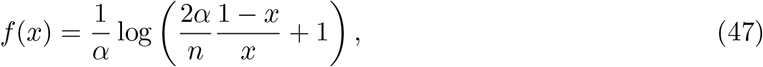

we approximate the expected coalescent interval as

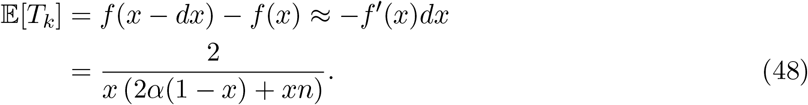

Note that while (48) is a large-*n* approximation, it allows that *α* might be of the same order of magnitude as *n*. Applying the same approximation to the combinatorial coefficient in (29) gives

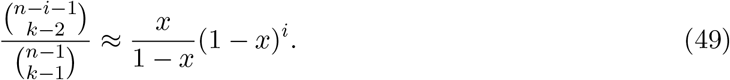

Finally, we approximate the sum in (46) with the integral

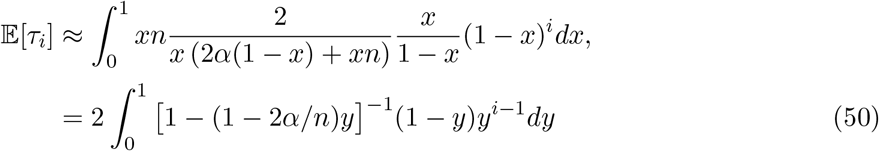

in which we changed variables (*y* = 1 − *x*) to make a connection to the hypergeometric function.

Thus we obtain

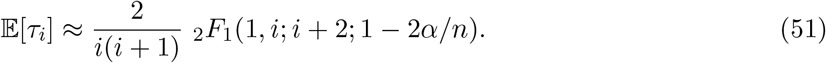

Equation (51) can be evaluated efficiently and the properties of the hypergeometric function are well know.

The suggested dependence of (51) on 1/*i*^2^, rather than the usual site-frequency prediction of 1/*i*, is consistent with the excess of low-frequency variants expected under population growth. As Slatkin and Hudson (1991) and others have observed, gene genealogies under very fast exponential growth are close to star trees. In this extreme, all variants will be singletons. From (51), when *α/n* is large, we have

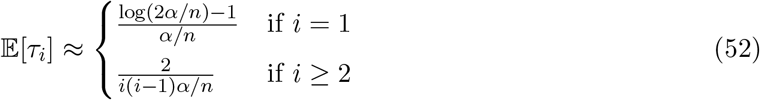

if terms of order (*α/n*)^−2^ or smaller are ignored. Thus singletons will dominate as expected when *α/n* is very large.

These results for exponentially growing populations, derived here using a coalescent approach, are identical in form to some results for “Luria-Delbrück distributions”, especially in application to cancer, derived using forward-time birth-death or branching processes (Luria and Delbrück, 1943; Lea and Coulson, 1949; Kessler and Levine, 2013; Ohtsuki and Innan, 2017; Gunnarsson et al., 2021; Cheek and Antal, 2018; Poon et al., 2021; Durrett, 2013, 2015). In particular, (50) has the same form as the approximation in equation (4) of Ohtsuki and Innan (2017) and as equation (33) in Gunnarsson et al. (2021). Equation (52) has the same form as the expression in Theorem 2 in Durrett (2013) if only the leading-order term is kept in (52) in the case *i* = 1.

Figure 4 shows the same quanties as Figure 2 but for the pure exponential growth model with *n* = 10^5^ and *α/n* = 3. The value *α/n* = 3 was chosen to roughly reproduce the ratio of singletons to doubletons observed for low-rate sites in the gnomAD data in Section 3.2. Figure 4A is directly comparable to Figure 2A, the only difference being whether 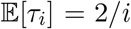 or comes from (51). As Figure 4A shows, recent rapid growth produces a single-mutation (*k*_1_ = 1, blue line) site-frequncy spectrum with an excess of rare variants and a deficit of common variants. So, compared to the constant-size case in Figure 2A, there is a diminished tendency to observe high-frequency variants when the number of latent mutations is larger, and a stronger tendency for the site-frequency count (*n*_1_) to be equal to or close to the number of latent mutations.

**Figure 4:**
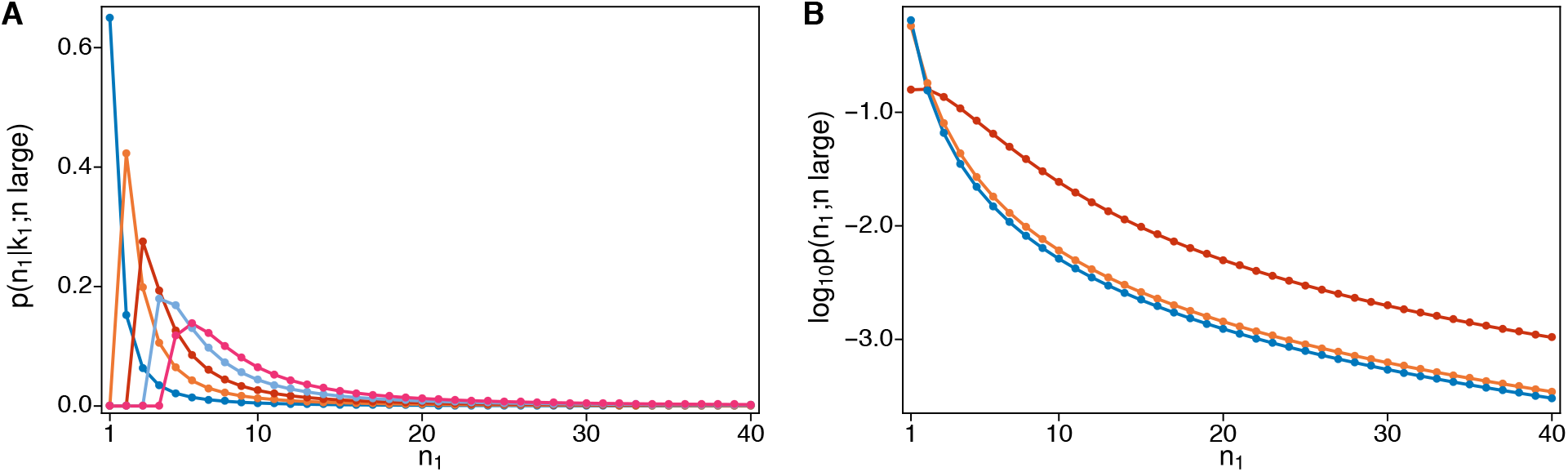
Plots of the same quantities shown in Figure 2 but for a sample of size *n* = 10^5^ under pure exponential growth with *α/n* = 3. Panel A: probability of observing *n*_1_ copies of allele 1 in the sample given *k*_1_ = 1, 2, 3, 4, 5 latent mutations. Panel B: log_10_-probability of observing *n*_1_ copies of allele 1 in the sample for three different mutation rates, corresponding to the values of *θπ*_1_: 0.002, 0.02 and 0.2 in Figure 2, but here expressed in terms of expected numbers of mutations on the gene genealogy (44): 0.024, 0.24 and 2.4. Probabilities in both panels are normalized to sum to one for *n*_1_ ∈ (1, 2, …, 40).

To make Figure 4B comparable to Figure 2B, we used (44) with *n* = 10^5^ and 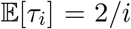 to compute the corresponding expected numbers of mutations on the gene genealogy for the three values of *θπ*_1_ in Figure 2 (0.002, 0.02, 0.2). The resulting expected numbers of mutations were 0.024, 0.24 and 2.4, the last being about equal to the value for the highest-rate sites in the gnomAD data in Section 3.2. We then computed *p*(*n*_1_; *n* large, *τ*) by averaging (34) over the distribution (33). Similar to Figure 2B, the two smaller values of the mutation rate give nearly indistinguishable results for the total count *n*_1_. But there is a dramatic difference for the largest mutation rate. In Figure 2B the prediction is distinctly L-shaped and thus similar to that for the lowest mutation rate, which again is 100-fold lower. In contrast, in Figure 4B singletons have a much lower chance of being observed. In fact, doubletons are slightly more likely than singletons. This relative excess of doubletons is due to the fact when there are two latent mutations these are highly likely to produce two copies of the variant under growth (Figure 4A) than under constant size (Figure 2A).

It is also of interest to know how the number of latent mutations in the ancestry of a rare variant depends on its count. Figure 5 depicts this for a series of increasing counts *n*_1_, from 1 to 16. Figure 5A shows the results for constant size, Figure 5B the corresponding results for pure exponential growth. The expected number of mutations on the gene genealogy is 2.4 in both cases. Regardless of the demography, if only one copy of the variant is observed, it must be due to one mutation. Otherwise, the results differ greatly for constant size versus growth. Under constant size, a variant observed multiple times in the sample can easily be due to a single mutation. Under growth, higher variant counts are more likely due to multiple mutations.

**Figure 5:**
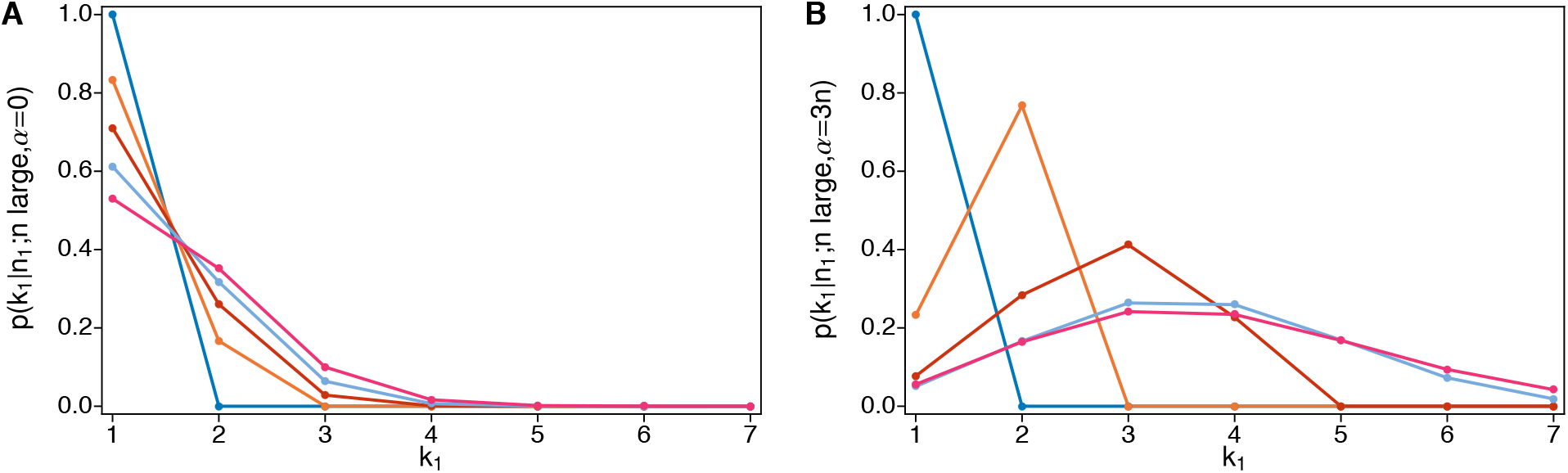
Probabilities of *k*_1_ = 1, 2, 3, 4, 5, 6, 7 latent mutations for increasing values of *n*_1_ — 1, 2, 4, 8, and 16, with blue for 1, orange for 2, and so on — when 2.4 mutations are expected on the gene genealogy of a sample of size *n* = 10^5^ (or equivalently *θπ*_1_ = 0.2 in the constant-size case). Panel A plots (25) with *θπ*_1_ = 0.2. Panel B shows the same probability computed using (33) and (34) under exponential growth with *α/n* = 3.

### 3.2 Application to human SNP data

We also used (33) and (34) to account for latent mutations in the ancestry of low-count variants in a subset of the gnomAD data (Karczewski et al., 2020). We took the approach described in the Supplementary Materials of Seplyarskiy et al. (2021), specifically obtaining estimates of relative branch lengths (43) from low-rate sites then using our new analytical result (34) to average over mutation counts. Rather than categorizing variants by trinucleotide context as in Seplyarskiy et al. (2021), we analyzed data from gnomAD version v2.1.1, presorted into 109 bins based on estimates of mutation rate by the method of Seplyarskiy et al. (2022, in prep.) which incorporates information from the six flanking bases on either side of a SNP, strand asymmetry, expression level, methylation promoter status. We did not use this information but our analysis assumes that variants within a bin have the same mutation rate.

The data consist of variant counts for synonymous mutations in the exomes of about 57K non-Finnish Europeans. Thus *n* ~ 114K although this varied by about 2% among sites because we required that sites were successfully genotyped in a minimum of 112K chromosomes. Importantly for our application, the data include monomorphic sites, i.e. sites with variant count equal to zero. gnomAD only provides *n* for polymorphic sites, so we imputed *n* for monomorphic sites using the nearest value at a polymorphic site within 100bp on either side of the focal site. After filtering for sequencing quality and coverage as well as removing mutation rate bins with fewer than 100 observed mutations, there are a total of 12, 338, 176 sites in 97 bins and 834, 486 of these are polymorphic.

Figure 6 shows the total numbers of sites (blue circles) and the number of monomorphic sites (orange circles) in each bin. The great majority of sites are in bins 1 through roughly 25. These have low mutation rates, as indicated by the close overlap of blue and orange circles, or nearly equal numbers of total sites and monomorphic sites. The widening gap between the blue and orange circles reflects the fact that higher-number bins have larger mutation rates. Estimates of the expected numbers of mutations per site for each bin range from 0.0097 for bin 1 to 2.23 for bin 97, with a mean of 0.083 (see below).

**Figure 6:**
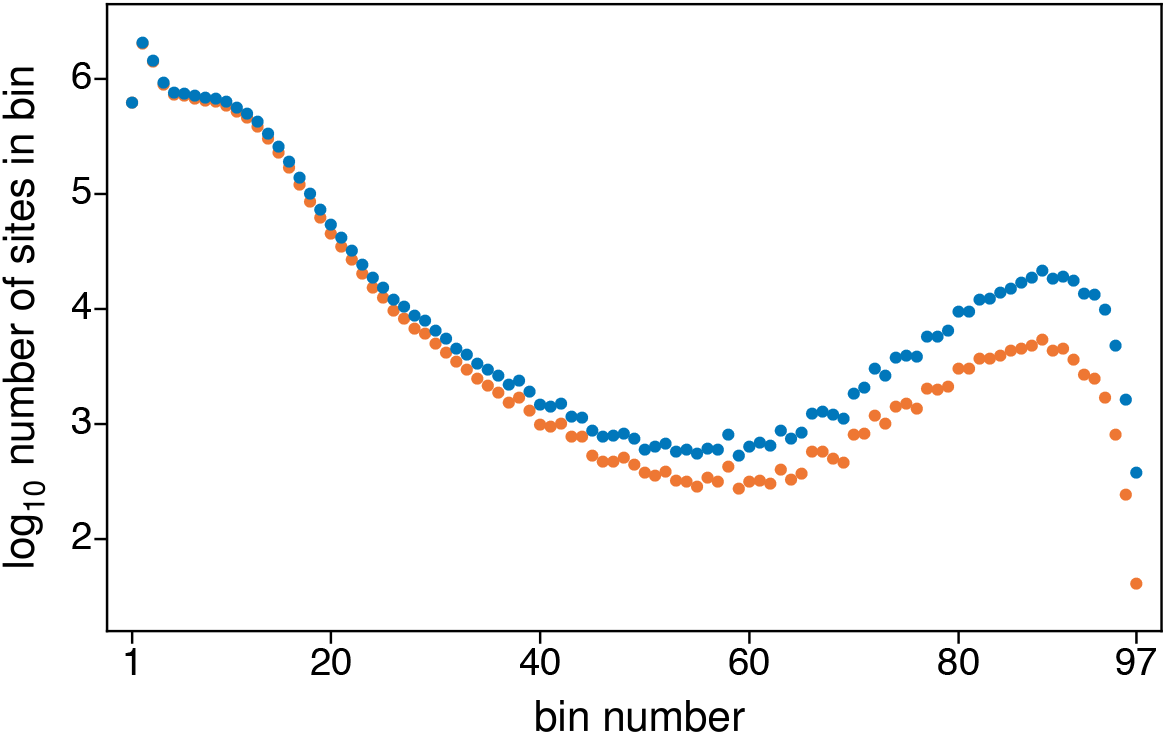
Blue circles show the total numbers of sites in the gnomAD data for each of the 97 bins. Orange circles show show the total numbers of monomorphic sites, i.e. sites with variant count equal to zero, in each bin. Lower mutation-rate bins are on the left, higher mutation-rate bins are on the right. The estimated mutation rates for bins 1 and 97 and about 9 times smaller and 27 times larger than the average (see text).

Each bin contains a mixture of different sequence contexts and different mutations. Again, we assume that within a bin these all have the same rate. We use *θπ*_1_ to denote this rate. Let *S_i_* be the number of sites in a given bin where *i* copies of a variant are observed in the sample. If a bin contains *L* total sites, then with reference to the notation in (2) we may write

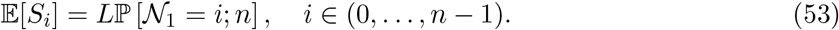

Thus we use *i* in place of *n*_1_ to avoid the additional subscript when we apply the results of the previous sections.

We compare observed and expected site-frequency counts for each bin based on an entirely empirical fit of our general results (33) and (34). This involves three steps. First, we use (33) and the proportion of monomorphic sites, *S*_0_/*L*, to estimate the total mutation rate in each bin, 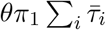, i.e as −log(*S*_0_/*L*). Next, based on (37), we use data for low-rate bins to estimate 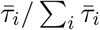 as *S_i_*/(*L* − *S*_0_) for *i* ∈ (1, …, 40). Finally, we compute the expectations 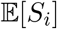, *i* ∈ (1, …, 40), for each bin using the estimated total mutation rate and the total number of sites for that bin. We do this using (33) and (34), and assuming that the 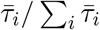 estimated from low-rate bins holds for all bins.

We used the combined variant counts for the first five bins to estimate the relative branch lenghts 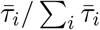. Our estimates of the total mutation rate for these bins range from 0.0097 to 0.037 with an average of 0.021. This is somewhat less than the smallest mutation rate in Figure 4 (also Figure 2) from which we can infer that these sites are unlikely to be affected by multiple mutations. Again, over all 97 bins, our estimates range from 0.0097 to 2.23, or about 230-fold from lowest to highest. The average across all bins is 0.083. Assuming that the latter corresponds to the genome average mutation rate per site, for which the usual estimate of *θ* from pairwise differences is about 1/1300 ~ 0.0077, we can infer that the expected number of mutations between a pair of (haploid) genomes is about 9 × 10^−5^ for the slowest sites and about 0.02 for the fastest sites.

Figure 7 compares the observed and expected variant counts, *S_i_* for *i* ∈ (1, …, 40), for bins 9, 50 and 92, chosen to represent a low-rate bin, a middle-rate bin and a high-rate bin. Figure 9 in the Appendix gives the plots for all 97 bins. Red circles show the observed counts. Blue lines trace the expected values. In making these plots, we grouped variant counts for which 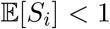. For bin 50 for example, this was true of variant counts *i* ∈ [12, 40] as noted in Figure 7B and again in the 50th panel of Figure 9. The values of “mutrate” displayed in these plots are the estimates of the expected number of mutations per site on the gene genealogy, i.e. the total mutation rate 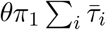, for each bin based on its proportion monomorphic sites.

**Figure 7:**
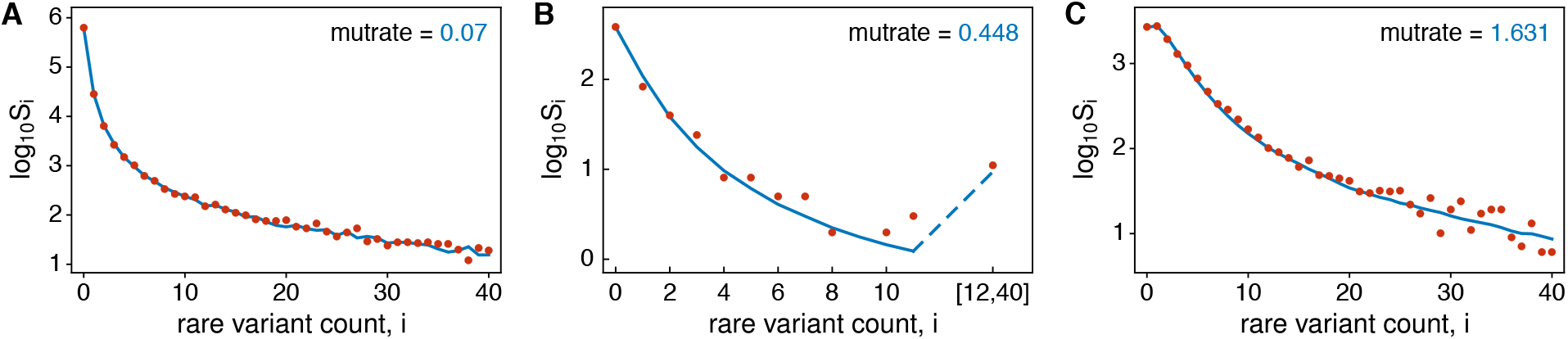
Examples of model fit for a (A) low-rate bin, (B) middle-rate bin, and (C) high-rate bin. Red dots are the data. Blue lines show expectations with “mutrate” 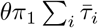 estimated using the new method. These are bins 9, 50 and 92 (cf. Figure 6).

The broad pattern from these plots is clear. For small total mutation rates (e.g. Figure 7A) the site-frequency spectrum is heavily weighted toward rare variants. For large total mutation rates (e.g. Figure 7C), i.e. when multiple latent mutations are likely, the site-frequency spectrum is shifted toward higher-frequency variants. As shown in Figure 6, the data contain fewer sites with intermediate mutation rates. In this case (e.g. Figure 7B), the site-frequency spectrum does show the expected intermediate pattern, but subject to considerable sampling error. Across the range of mutation rates, the empirical model, which uses low-rate sites to estimate relative branch lengths 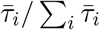 and assumes these hold for all sites, fits the data well.

As can be seen in Figure 7A and the first 20 or so panels of Figure 9, the empirical estimates of 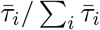 include fluctuations due to sampling error for higher-count variants. The combined data for the first five bins have *S_i_* ranging from 71 to 38 for *i* ∈ [30, 40]. The presence of these fluctuations helps illustrate a subtler phenomenon, namely the smoothing which occurs at larger mutation rates (e.g. Figure 7C). For reference, the combined data for the first five bins have *S_i_* in the thousands for the low-count variants. From these, the estimated chance that a latent mutation is a singleton is about 64%, followed by 13% for doubletons and 6% for tripletons. By comparison, the chance is less than 0.1% for each variant with count *i* ∈ [25, 40]. The predictions 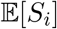 are smoothed for higher-count variants at larger mutation rates because they are mixtures. For example, two latent mutations will come in counts 1 and *i* − 1, 2 and *i* − 2, or 3 and *i* − 3 with approximate relative proportions 64:13:6.

## 4 Discussion

In this work, we modeled the mutational ancestry of a rare variant in a large sample. Under the standard neutral model of population genetics with *K*-allele parent-independent mutation, we found that co-segregating rare variants may be treated independently and that the Ewens sampling formula gives the probabilistic structure of latent mutations in their ancestries. We obtained more general results, which hold under changing population size, by modeling latent mutations as independent Poisson random variates.

Our aim has been to describe how the site-frequency spectra of rare variants in large samples are affected recurrent mutation. The key parameters for a variant in count *i* turn out to be its expected total rate of mutation on the gene genealogy of the sample (here denoted 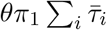) and the expected relative lengths of branches in the gene genealogy which have *i* descendants 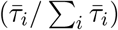. Under the standard neutral model 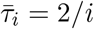.

We obtained new results for 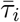 under exponential population growth and used these to illustrate how recurrent mutation affects the site-frequency spectrum differently than under constant size. Lastly, we showed that our general results provide a good fit to synonymous variation among a large number of (non-Finish European) individuals in the human Genome Aggregation Database (Karczewski et al., 2020), suggesting that, whatever the causes of deviations from 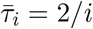 might be for this sample, differences in mutation rate can explain differences in site-frequency spectra among different kinds of sites.

Our application was empirical. We did not fit a demographic model, but following Seplyarskiy et al. (2021) used low-mutation-rate sites to estimate relative branch lengths and assumed these hold for all sites. Site-frequency spectra are a rich source of information about population-genetic phenomena but are of somewhat limited use in disentangling their effects (Myers et al., 2008; Bhaskar and Song, 2014; Terhorst and Song, 2015; Lapierre et al., 2017; Rosen et al., 2018). When low-mutation-rate sites are plentiful enough to provide stable estimates of relative branch lengths, this empirical method offers a way to control for myraid factors and isolate the effects of variation in mutation rate.

We began with a *K*-allele model with parent-independent mutation, and used its sampling probabilities in our computations for constant-size populations. We conjecture that our findings will hold for general mutation models because conditioning on a rare variant in a large sample means that the ancestral allele will be the common allele with very high probability. Then the relevant mutation rate in any model will be the rate of the production of the rare allele from the common allele.

We have described our general results as being for populations which may have changed in size. This is appropriate for the general coalescent model of Griffiths and Tavaré (1998) which we used to portray our results and assumed in the proofs in the Appendix. Strictly speaking, though, the general coalescent does not require a generative model for the times between coalescent events, *T_k_* for *k* ∈ (2, …, *n*). Selection might be part of the reason they differ from the predictions of the standard neutral coalescent. This may be true, for example, for the synonymous exome data from gnomAD we analyzed.

In fact, the derivation of (33) and (34), with associated results from (30) to (38), does not even require interpretation in terms of coalescence times. These equations hold just as well if we replace 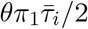 with an arbitrary rate parameter *λ_i_* for the production of mutants in count *i*, potentially of an allele which is under selection. The case of a fixed tree, with fixed *τ_i_* not from a generative model, considered in the Appendix is an example. The modified Poisson Random Field model of Desai and Plotkin (2008) is another, in which *λ_i_* was the rate under additive selection (and the independent-Poissons assumption was applied to all counts in the sample). We have shown in detail how our results follow from the standard neutral coalescent or diffusion model and its extension the general coalescent model. As with our conjecture about general mutation models, we expect these results can be applied to latent mutations of alleles under various kinds of selection and a range of demographies (Lange and Fan, 1997; Dorman et al., 2004; Lambert, 2011; Kaj and Mugal, 2016; Torres et al., 2020; Müller et al., 2022) because they are for rare variants in large samples.

## Acknowledgments

We thank Vladimir Seplyarskiy for helpful discussions and assistance with the mutation-rate binned SFS data from gnomAD. This work was supported in part by National Science Foundation grants DMS-1855417 and DMS-2152103, and Office of Naval Research grant N00014-20-1-2411 to Wai-Tong (Louis) Fan; and by National Institutes of Health grants R35-GM127131, R01-MH101244, U01-HG012009 and R01-HG010372 to Shamil Sunyaev.

## A Appendix

## A.1 Time-dependent conditional ancestral process

Here we study the conditional ancestral process in detail and provide the justification for (22) and (23).

Let 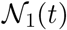 and 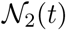 be the numbers of rare alleles and common alleles respectively at time *t*. From (19), (20) and (21), the stochastic process 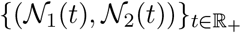 is a continuous-time Markov chain on 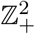 with total rate of events 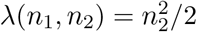 and one-step transitions

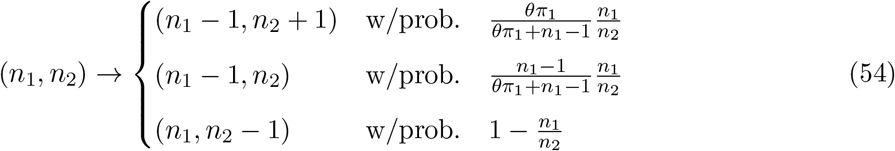

Let ℙ_**n**_ be the probability measure for this process starting at **n** = (*n*_1_, *n*_2_), and define the random times

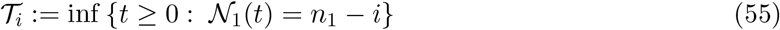

to be the times at which the first coordinate of the process decreases to *n*_1_ − *i* for 1 ≤ *i* ≤ *n*_1_, with 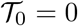. We have 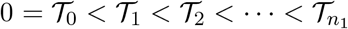 almost surely under ℙ_**n**_, and the process 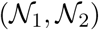 visits the following points in order 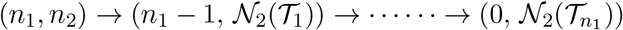.

In Theorem 1 we describe the joint distribution of the hitting times 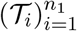 and the locations 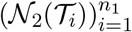 as *n*_2_ → ∞.

### Theorem 1

*As n*_2_ → ∞, *the random vector*

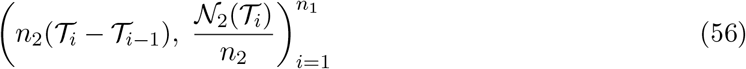

*in* 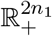 *converges in distribution under* ℙ_**n**_ *to the random vector*

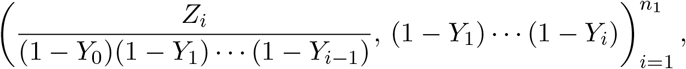

*where Y*_0_ = 0, *and* 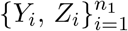 *are independent random variables with probability density functions*

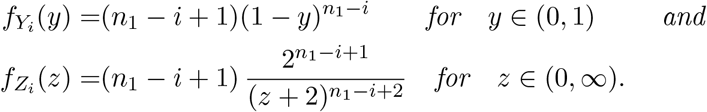

**Remark 1** (Mean of 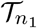). Note that

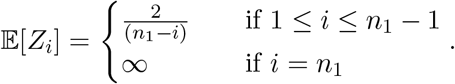

Hence for *n*_1_ ≥ 2, Theorem 1 implies that 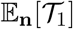 is of order 1/*n*_2_ and gives the second part of (23) in the main text. In contrast, when *n*_1_ = 1, 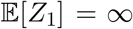 and 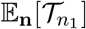 is no longer of order 1/*n*_2_. Indeed, when *n*_1_ = 1, 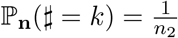 for *k* ∈ {0, 1, · · · *n*_2_ − 1} by (59). Hence by (57) and Fubinni’s theorem,

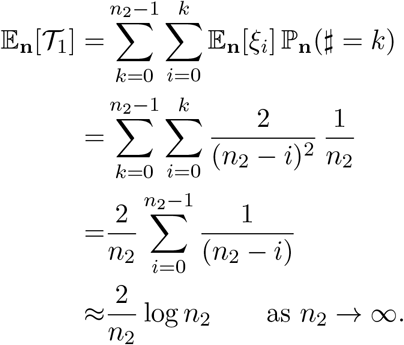

These give (22) in the main text.

**Remark 2** (Mean of 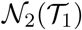). By (58) and Theorem 1,

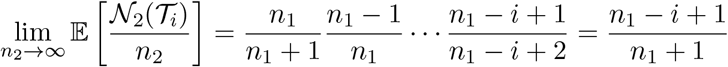

for 1 ≤ *i* ≤ *n*_1_. This gives (23) in the main text.

## A.1.1 Proof of Theorem 1

To explain the key idea we first establish weak convergence of 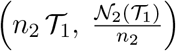, i.e. of the marginal distribution for *i* = 1 in (56). By definition, 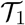 is given by

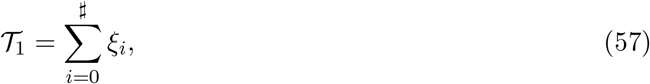

where *#* is the number of downward jumps in second coordinate of the process starting at (*n*_1_, *n*_2_) up to the first decrease in the first coordinate. The variables 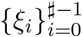 are the times between these downward jumps, with *ξ_#_* being the time to the final jump starting at (*n*_1_, *n*_2_ − *#*). This last jump is the one which decreases the first coordinate. Observe that 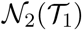 is either *n*_2_ − *#* or *n*_2_ − *#* + 1. Given *#*, 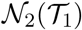 is equal to

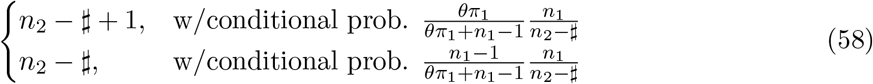

which correspond to a non-empty mutation event and a coalescent event of type 1 respectively. These follow from (54).

The probability mass function of *#* is given by 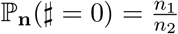 and, for *k* ∈ {1, 2, · · ·, *n*_2_ − *n*_1_},

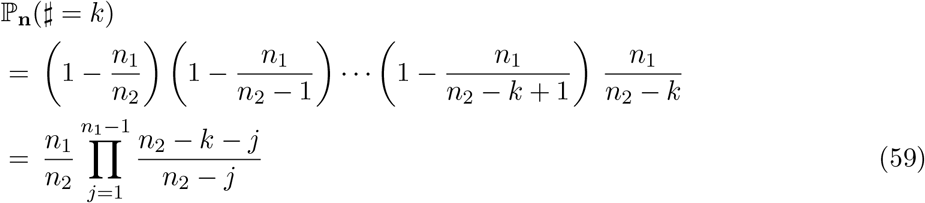

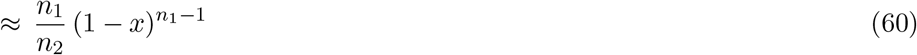

as *n*_2_ → ∞ and 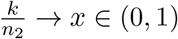. Hence ℙ_**n**_(*#* > *n*_2_ − *n*_1_) = 0 and, for *k* ∈ {0, 1, 2, · · ·, *n*_2_ − *n*_1_ − 1},

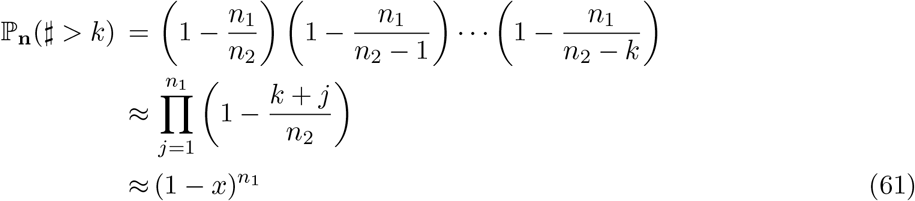

as *n*_2_ → ∞ and 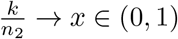.

### Lemma 1

*As n*_2_ → ∞, *we have convergence in distribution*

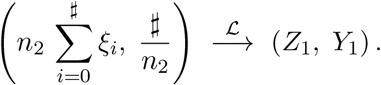

*with Z*_1_ *and Y*_1_ *as defined in Theorem 1*.

**Proof of Lemma 1.** It suffices to show that

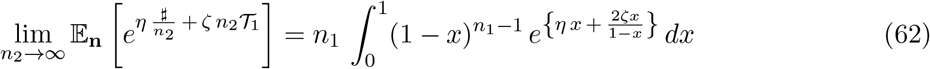

for *η* ∈ ℝ and *ζ* ∈ (−∞, 0]. Since *ξ_i_* ~ Exp(*λ*(*n*_1_, *n*_2_ − *i*)),

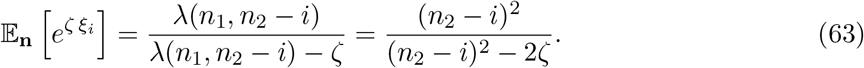

By (57), (59) and (63),

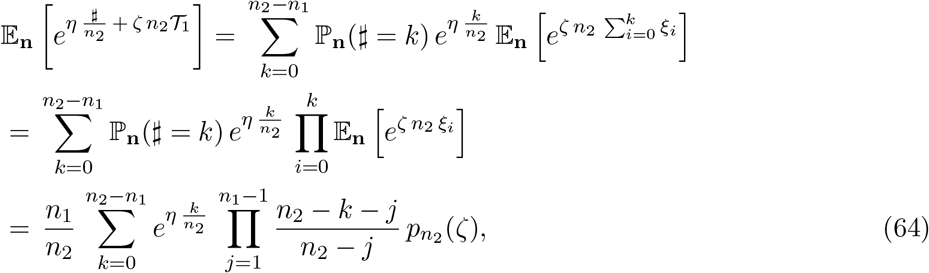

where

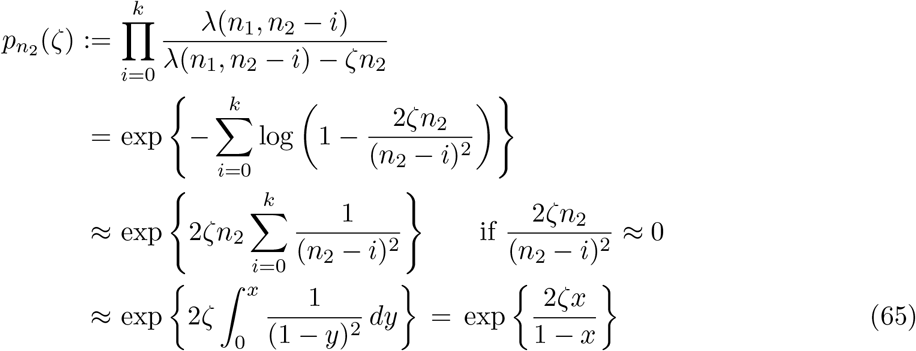

if 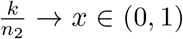 and *n*_2_ → ∞. Putting (65) and (60) into (64), we obtain the desired (62) and thus Lemma 1.

We now return to the proof of Theorem 1. Lemma 1 implies that 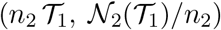 converges in distribution to (*Z*_1_, 1 − *Y*_1_) as *n*_2_ → ∞. Since *Y*_1_ < 1 almost surely, we have 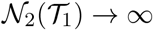 in the sense that

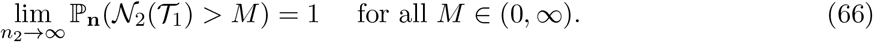

As in (57), by definition, 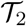 is given by

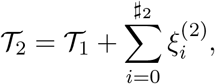

where *#*_2_ is the number of downward jumps starting in state 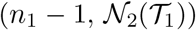 up to the second decrease in the first coordinate, i.e. to *n*_1_ − 2. Like before, 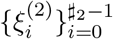 are the times between these jumps, with 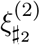 being the time for first coordinate to hit *n*_1_ − 2 starting at the penultimate states 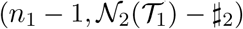. As in (58), 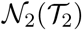 is either 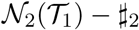 or 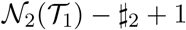.

As *n*_2_ → ∞, 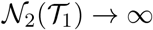 in the sense of (66). Hence the same argument that leads to Lemma 1 can be applied again, starting at the new location 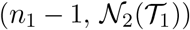. More precisely, by computing moment generating functions as before, and applying the strong Markov property of the random walk 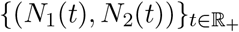 at the stopping time 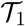, we obtain the joint convergence

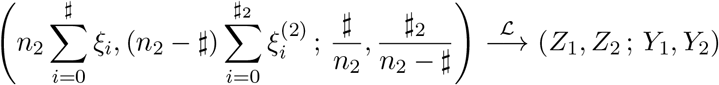

under ℙ_**n**_ as *n*_2_ → ∞, where {*Z*_1_, *Z*_2_, *Y*_1_, *Y*_2_} are independent variables defined in Theorem 1. This implies the convergence in distribution

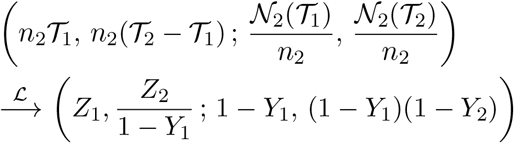

under ℙ_**n**_, as *n*_2_ → ∞. Continuing this way, by letting *#_i_* be the number of downward jumps starting at 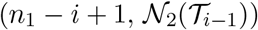 before hitting the vertical line {(*n*_1_ − *i*, *y*) : *y* ∈ ℤ_+_} for *i* ≥ 1, we obtain the desired convergence in Theorem 1.

## A.2 Low-count branches of general coalescent trees

Here we prove the non-nestedness and Poisson-independence of low-count mutations, which we assumed in Section 2. We do this first for fixed trees then for the random, general coalescent trees of Griffiths and Tavaré (1998). We also present the computation of the probability generating function, *G_n,k_*(*x, y*), of the count of the variant of interest and its number of latent mutations.

## A.2.1 Nested mutation on a fixed tree

Let **T**_*n*_ be a fixed (non random) tree with *n* leaves. We suppose the tree is ultrametric, that is the leaves have the same distance *H_n_* from the root. We call *H_n_* the height of **T**_*n*_. Consistent with the main text, we adopt the following notation for some relevant properties of **T**_*n*_, for the most part suppressing the dependence on *n* for simplicity:

1. *T_k_* is the length of the time during which there are exactly *k* lineages ancestral to the sample, for *k* ∈ {2, 3, · · ·, *n*}.
2. *τ_j_* for *j* ∈ {1, · · ·, *n* − 1}, is the total length of branches in **T**_*n*_ that have *j* descendants. We suppose there are *m_j_* such branches with lengths 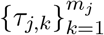. Then 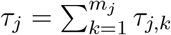.
3. *T*_total_ is the total branch length, the sum of all the branches in **T**_*n*_, which is equal to 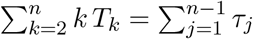.
4. For a positive integer *b*, we define a collection 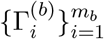 of disjoint connected subtrees of the coalescent tree as follows: Each of the *m_b_* branches with *b* descendants in the sample (say the *i*-th one) subtends *b* leaves in the coalescent tree and gives rise to a subtree 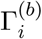 which contains that branch. We say **nested mutation up to count** *b* occurs on **T**_*n*_ if there exist two mutations on 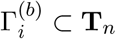 for some *i* ∈ {1, 2, · · ·, *m_b_*}. Figure 8 illustrates this for *b* = 4.

We assume that mutations arise as a Poisson point process on the tree with constant rate *θ*/2 per unit length. Theorem 2 below holds for any fixed ultrametric tree (it can be binary or have multiple mergers, or even be a star tree).

**Figure 8:**
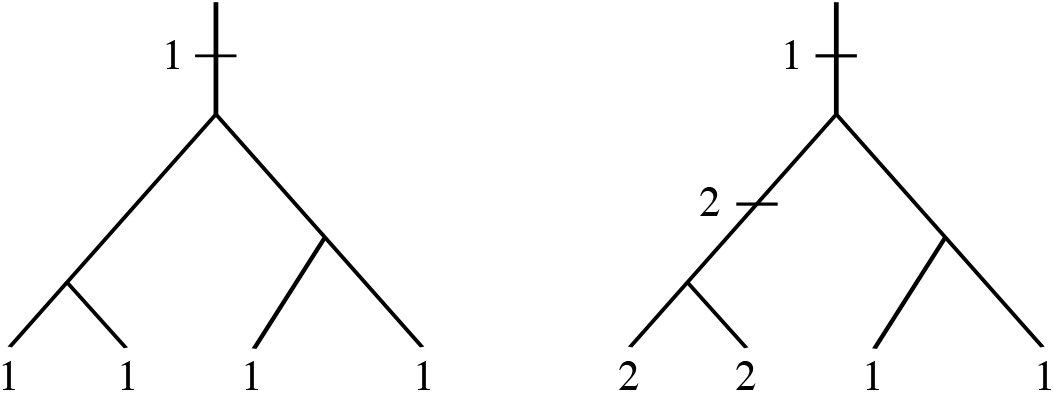
Two subtrees in 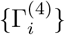. The subtree on the left has one mutation which is labeled 1 and has count four. The subtree on the right has nested mutations, with the mutation labeled 1 in count two and another labeled 2 also in count two.

**Figure 9:**
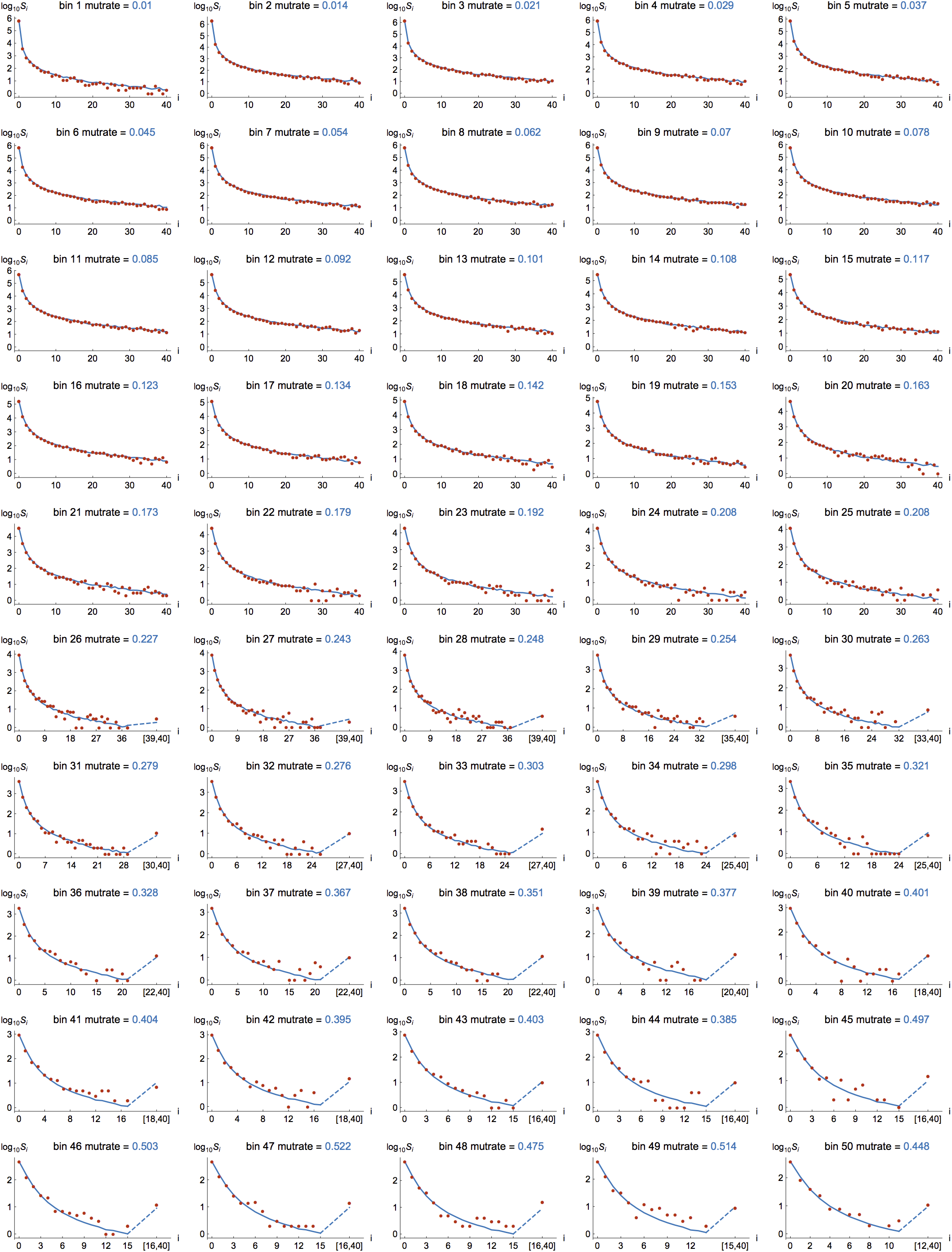
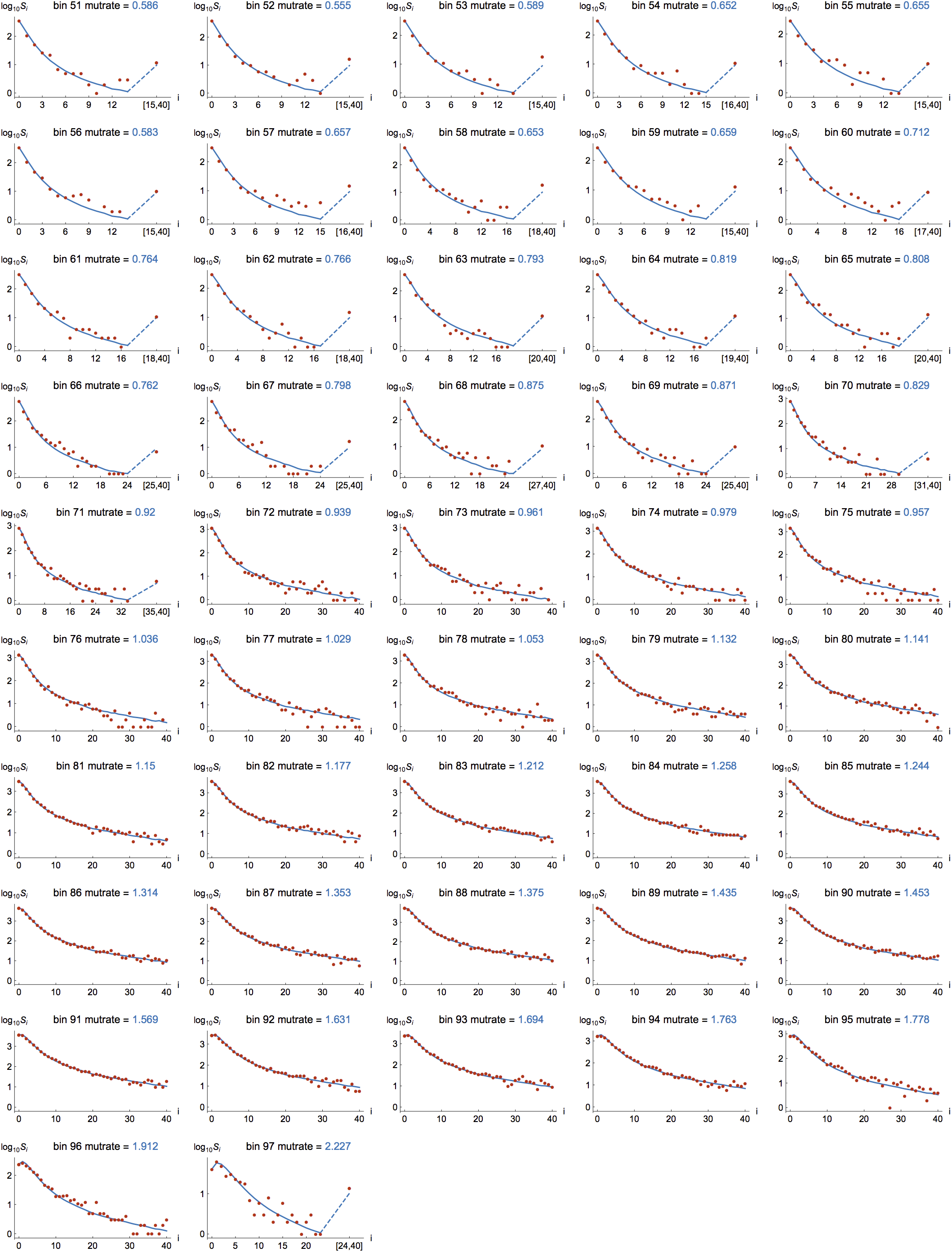
Plots like those in Figure 7 for each of the 97 mutation-rate bins.

### Theorem 2 (Nested mutation on fixed trees)

*Let* **T**_*n*_ *be a fixed ultrametric tree with n leaves. For any positive integer b and for any θ* ∈ (0, ∞), *the probability that nested mutation up to count b occurs is bounded above by*

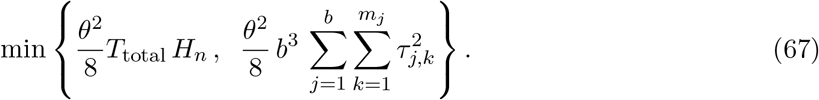

*In particular, the probability that nested mutation up to count b occurs tends to 0, as n* → ∞, *if* 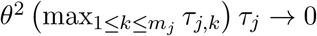 for 1 ≤ *j* ≤ *b*.

**Remark 3**. There is good evidence that the upper bound bound 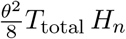 is actually small for humans. For the gnomAD data we analyze in the main text, the expected number of mutations per site (*θT*_total_/2) is between about 0.009 and 2.13. So *θT*_total_/2 is not big with high probability. The rest of the upper bound, *θH_n_*/4, should be proportional to the average pairwise difference per site (very nearly equal to this for random Kingman coalescent trees and large *n*) and this ranges from about 9 × 10^−5^ to about 0.02 for these same data. See Section 3.2.

**Remark 4**. The simpler bound 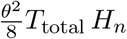 can be weaker than the other bound 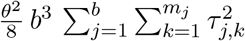 in (67) for large *n*. For the Kingman coalescent, 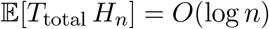 is larger than 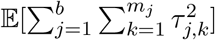 since the latter tends to 0 as *n* → ∞, by (70). For a star tree, however, both bounds are approximately 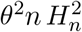 (up to a multiplicative constant).

*Proof*. The total number *M_n_* of mutations on **T**_*n*_ is a Poisson variable with mean 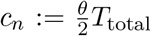. Given the tree **T**_*n*_ and *M_n_* = *k*, the *k* mutations are uniformly distributed on the tree. Hence the conditional probability that two given mutations are on the same subtree 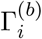 for some *i* is equal to

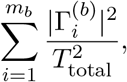

where 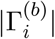 is the total branch lengths of the subtree 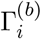. Since there are *k*(*k* − 1)/2 ways to choose two mutations out of *k*,

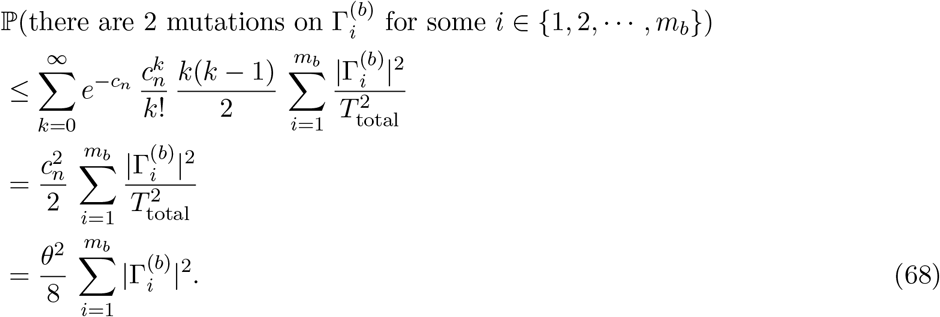

From here we can apply the simple bound 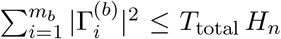 to obtain the first bound 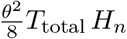 in (67). To get the second bound in (67), note that 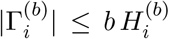 for all 1 ≤ *i* ≤ *m_b_*, where 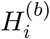 is the height of the subtree 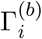.

Furthermore, 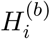 is the sum of at most *b* branch lengths, one from {*τ_j,k_*} for *j* = *b*, *b* − 1, · · ·, 2, 1, and these branches are pairwise disjoint for different *i*’s (for 1 ≤ *i* ≤ *m_b_*). Hence

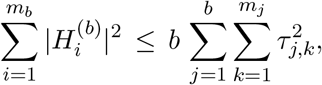

where we used the general inequality 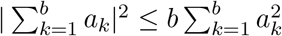. The bound in (67) now follows by putting these into (68).

A mutation on a tree (called a latent mutation in the main text) is said to **have count** *j* if the mutation is the most recent mutation in the lineages of exactly *j* individuals at the leaves of the tree; see Figure 8.

### Theorem 3 (Poisson approximation for counts on a fixed tree)

*Let* **T**_*n*_ *be a fixed coalescent tree with n leaves for n* ≥ 2. *Let a_j_ be the number of mutations on* **T**_*n*_ *with counts j. If the probability that nested mutation up to count b occurs tends to 0 as n* → ∞, *then for any positive integer b and any θ* ∈ (0, ∞), *the variables* 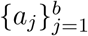 *are asymptotically independent and* 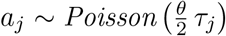 *for* 1 ≤ *j* ≤ *b*.

*Proof*. If there is no nested mutation up to count *b*, then *a_j_* is also equal to the number of mutations on the branches in **T**_*n*_ that have *j* descendants, for 1 ≤ *j* ≤ *b*. Since these branches have total length *τ_j_* and they are disjoint for different *j*’s, the result follows from the assumption that mutations occur as a Poisson point process on the tree **T**_*n*_ with rate *θ*/2.

## A.2.2 Nested mutation on random trees

We now suppose the tree **T**_*n*_ is a *random* binary tree (for *n* ≥ 2), in particular the general coalescent tree of Griffiths and Tavaré (1998). For each *n* ≥ 2, 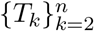 is a sequence of positive random variables representing the times during which there are *k* lineages in **T**_*n*_. The branching structure of **T**_*n*_ is independent of the times 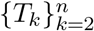. Looking forward in time, whenever there is a branching event, an existing lineage is chosen uniformly at random to split into two.

Following Griffiths and Tavaré (1998, eqn. (2.2)) we let λ(*t*) be the the population size at time *t* in the past divided by the current population size. As in (45), λ(*t*) = *e*^−*αt*^ with *α* > 0 corresponds to an exponentially growing population.

### Theorem 4 (Nested mutation on random trees for fixed *θ*)

*Let b* ∈ ℕ. *Suppose for* 1 ≤ *j* ≤ *b*,

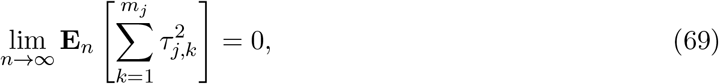

*where the expectation* **E**_*n*_ *averages over all realizations of* **T**_*n*_. *Then the probability that nested mutation up to count b occurs is bounded above by C_b,n_ θ*^2^, *where* {*C_b,n_*}_*n*≥2_ *are constants that tend to 0 as n* → ∞. *Furthermore*, (69) *holds for the generalized coalescent trees of Griffiths and Tavaré (1998) when* sup_*t*≥0_ λ(*t*) < ∞ *(which includes any growing population)*.

*Proof*. The first statement follows directly from Theorem 2. By the fact 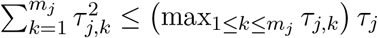 and the Cauchy-Schwarz inequality, we have

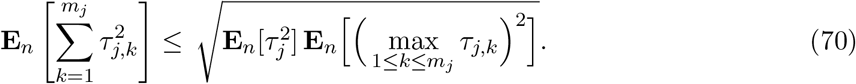

Hence assumption (69) is satisfied if

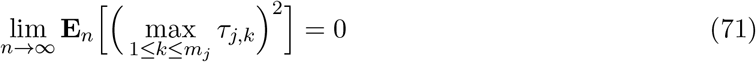

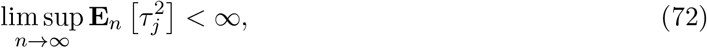

for 1 ≤ *j* ≤ *b*. The second statement now follows from Lemma 2, Lemma 3, and Proposition 1 below.

Lemma 2 concerns assumption (71). For reference, we note that it is satisfied, and hence (71) is satisfied, if *T_k_* are exponential variables with parameter *λ_k_* where 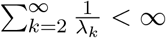. This is true for the Kingman coalescent which has *λ_k_* = *k*(*k* − 1)/2.

### Lemma 2

*Suppose* 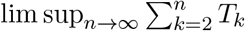 *has finite p-th moment, where p* > 0. *Then* 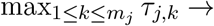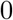 *in L^p^, as n* → ∞.

*Proof*. Consider the random tree **T**_*n*_ and recall that *T_k_* is the length of the time during which there are exactly *k* lineages ancestral to the sample in **T**_*n*_. These *k* lineages are segments of length *T_k_* of the branches of the genealogy, and each of them is called a line of state *k*.

Let 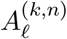 be the number of descendants in **T**_*n*_ of the *ℓ*-th line of state *k*. Note that 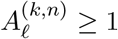 for *ℓ* ∈ {1, 2, · · ·, *k*}, and 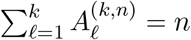. Since the branching structure is independent of 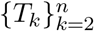, we can assume without loss of generality that 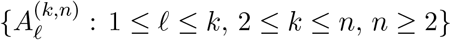 are all defined on the same probability space. By exchangeability—in particular see Bertoin (2006, Proposition 2.8)—the random vector 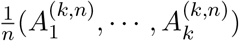 converges almost surely to a random vector that has the symmetric Dirichlet distribution on the simplex 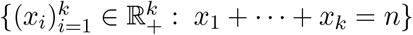. Therefore, with probability one,

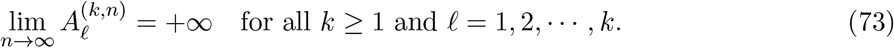

Since 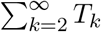 is finite almost surely, the trees {**T**_*n*_}_*n*≥2_ have uniformly bounded height almost surely. So (73) implies that with probability one,

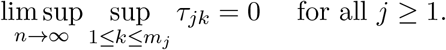

Since 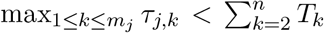, by the assumption on {*T_k_*} and the Dominated Convergence Theorem, 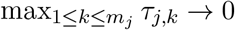 in *L^p^* as *n* → ∞.

Next consider assumption (72). For the Kingman coalescent, *τ_j_* is close to its mean **E**_*n*_[*τ_j_*] = 2/*j* because for *n* large enough,

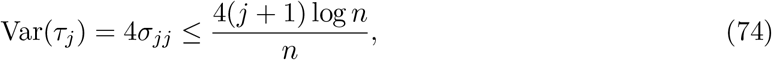

where *σ_jj_* is defined in Fu (1995, eqns. (1)–(2)). This follows from the fact 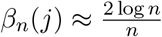 as *n* → ∞ for each *j* ≥ 1 (Fu, 1995, eqn. (5)). Hence

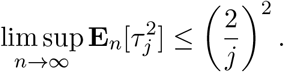

### Lemma 3

*Suppose there exists a constant C*_*_ ∈ (0, ∞) *such that*

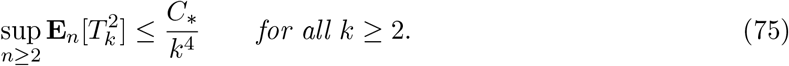

*Then* 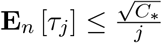 *for all j* ≥ 1 *and* 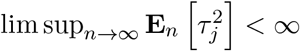.

*Proof*. For realized values of *T_k_*, the argument in Fu (1995, page 181) gives

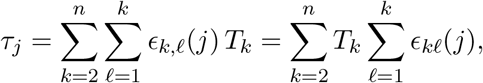

where 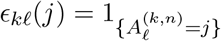 is the indicator variable, where 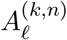 is the number of descendants in **T**_*n*_ of the *ℓ*-th line of state *k* defined in the proof of Lemma 2.

Using the independence between {*T_k_*}_*k*≥2_ and the branching structure, and following the notation in Fu (1995, eqns. (18)–(19)), the conditional expectation of *τ_j_*, given 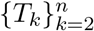, is

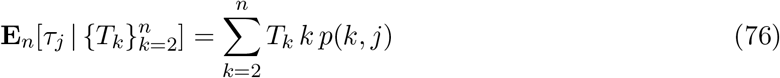

and that of 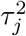, given 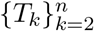, is

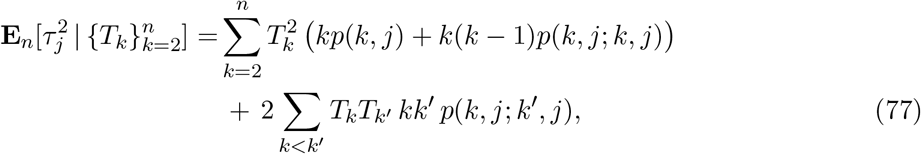

where the deterministic functions *p*(*k*, *j*), *p*(*k*, *j*; *k*′, *j*) do not depend on {*T_k_*}. From Fu (1995),

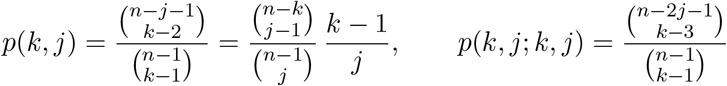

and for 2 ≤ *k* < *k*′ ≤ *n*,

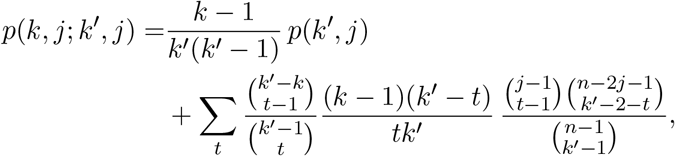

where the sum is taken over 1 ≤ *t* ≤ min{*j*, *k*′ − 2, *k*′ − *k* + 1}.

The first and the second moments of *τ_i_* are obtained averaging over 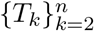 in (76) and (77). The bound 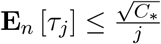 follows from the same calculation in Fu (1995, eqn. (22)). By (77), the fact 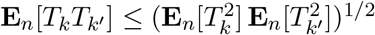 and assumption (75), 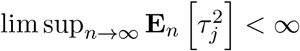 holds also for our random trees.

**Remark 5**. As in Theorem 2, we can use an alternate assumption than 69. For any positive integer *b*, the probability that nested mutation up to count *b* occurs is bounded above by 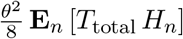 which tends to 0 if *θ*^2^ **E**_*n*_ [*T*_total_ *H_n_*] → 0. For Kingman coalescent trees, this would require that *θ* → 0.

We now check that the assumption (69) in Theorem 4 holds for the generalized coalescent tree of Griffiths and Tavaré (1998).

### Proposition 1

*Suppose C*_0_ ≔ sup_*t*≥0_ λ(*t*) < ∞. *Then* {*T_k_* : 2 ≤ *k* ≤ *n*, *n* ≥ 2} *satisfy the conditions in both Lemma 2 (with p* = 2*) and Lemma 3. In particular*, (69) *is satisfied and so the conclusion of Theorem 4 holds*.

*Proof*. The joint distribution of 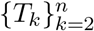 is determined by the function λ; see Griffiths and Tavaré (1994b). We can construct 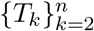 in terms of λ as follows: let 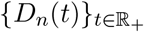 be a pure death process with rate 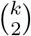 at state *k* ∈ {1, 2, · · ·, *n*}, starting at *D_n_*(0) = *n*, and let

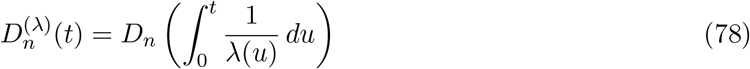

be a time-changed pure death process. Then

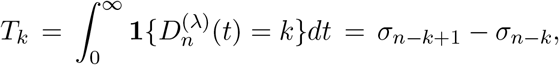

for 2 ≤ *k* ≤ *n*, where *σ*_1_ < *σ*_2_ < · · · < *σ*_*n*−1_ are the jump times of 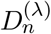 (by convention *σ*_0_ = 0).

By (78), the jump times of the pure death process *D_n_*, denoted by 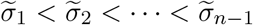, are given by 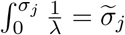 for 1 ≤ *j* ≤ *n* − 1. Hence, with the convention *σ*_0_ = 0, for 0 ≤ *j* ≤ *n* − 2 we have

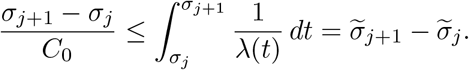

These give 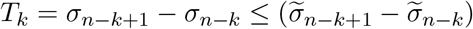 for all 2 ≤ *k* ≤ *n*.

Since 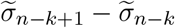 is equal in distribution to the analogue of *T_k_* for the Kingman coalescent, *T_k_* is stochastically dominated by *C*_0_ times an exponential variable with parameter *k*(*k* − 1)/2 for all 2 ≤ *k* ≤ *n*. The desired statement now follows since (71) and (72) are satisfied.

## A.2.3 Replacing *τ_j_* by its mean

By using the expected coalescence times denoted 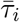 in the main tex, we implicitly assumed that different sites have different trees and that these are all drawn from the same distribution. Theorem 5 below asserts that even though the mutant counts at each site are conditional on the realization of the tree at that site, we can replace *τ_j_* by its expectation **E**_*n*_[*τ_j_*] in Theorem 3 when the trees are random and satisfy suitable assumptions. The key reason is that *τ_j_* is close to its mean, as made precise in Lemma 4.

### Lemma 4

*Suppose* (75) *holds and that the covariance*

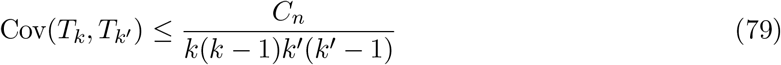

*for* 2 ≤ *k* < *k*′ ≤ *n and n* ≥ 2, *where* {*C_n_*} *is a sequence that tends to 0 as n* → ∞. *Then for each j* ≥ 1, *the variance* Var(*τ_j_*) → 0 *as n* → ∞. *In particular*, 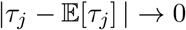 *in L*^2^(ℙ) *as n* → ∞.

*Proof*. By further taking expectations in (76) and (77) with respect to **E**_*n*_, we obtain the variance

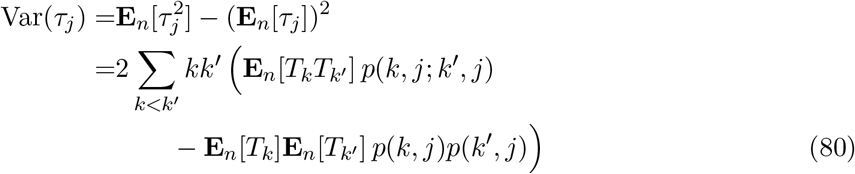

up to an 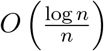 term. This follows from Fu (1995, eqns. (24)–(25)) and assumption (75) in Lemma 3. This also leads to (74).

By assumptions (75) and (79), the double sum in (80) is bounded above by

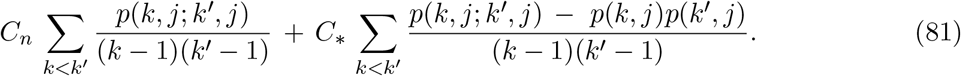

By Fu (1995, eqns. (29) and (22)), the first and second terms of (81) are of order *o*(*n*) and 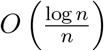 respectively, as *n* → ∞ for each *j* ≥ 1. The completes the proof of lim_*n*→∞_ Var(*τ_j_*) = 0. The latter implies, by Chebyshev’s inequality, that 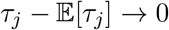 in *L*^2^ as *n* → ∞.

### Theorem 5 (Poisson approximation for counts across loci)

*Let* {**T**_*n*_}_*n*≥2_ *be a sequence of random coalescent trees which are the generalized coalescent trees of Griffiths and Tavaré (1998). Suppose* sup_*t*≥0_ λ(*t*) < ∞ *and assumption* (79) *holds. Let a_j_ be the number of mutations on* **T**_*n*_ *with counts j. Then for any positive integer b and any θ* ∈ (0, ∞), *the variables* 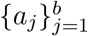 *are asymptotically independent and* 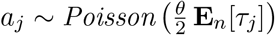 *for* 1 ≤ *j* ≤ *b, as n* → ∞.

*Proof*. By Theorem 4, the probability that nested mutation up to count *b* occurs tends to 0 as *n* → ∞. The result then follows from Lemma 4 and Theorem 3.

It can be checked that exponentially growing popolations clearly satisfy sup_*t*≥0_ λ(*t*) < ∞ and also assumption (79). The conclusions of Theorems 4 and 5 then hold for the generalized coalescent trees of Griffiths and Tavaré (1998) when λ(*t*) = *e^αt^* for *t* ∈ ℝ_+_ for some *α* > 0.

Equipped with Theorem 5, we write 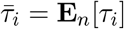 as in the main text and compute the probability generating function *G_n,k_* of the count of the variant of interest and its number of latent mutations. The count of the variant of interest is *k* = Σ*_i_ ia_i_* and its number of latent mutations is *k* = Σ*_i_ a_i_*. Hence

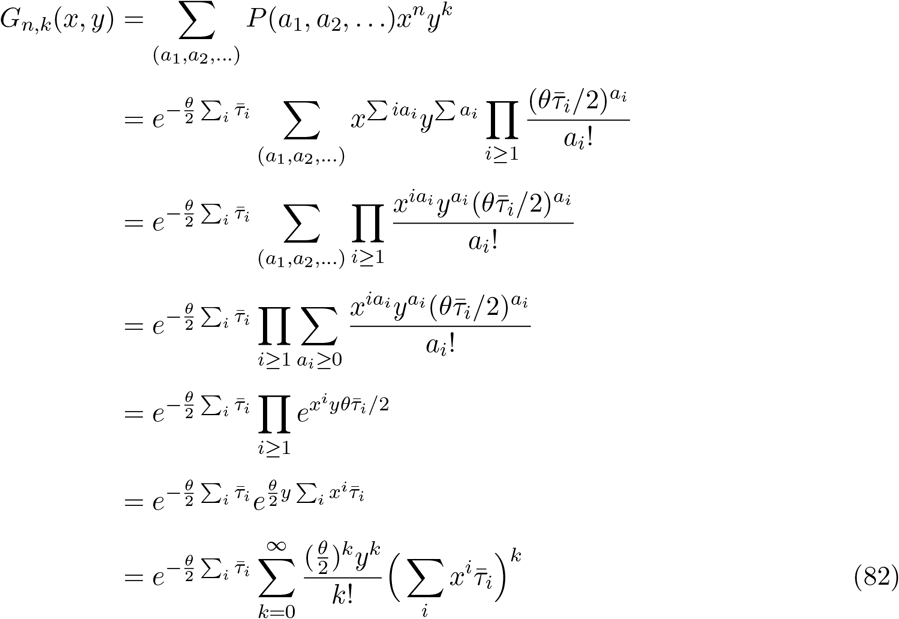

as declared in the main text.

